# Tissue-intrinsic γδ T cells critically regulate Tissue-Resident Memory CD8 T cells

**DOI:** 10.1101/2022.01.19.476598

**Authors:** Miguel Muñoz-Ruiz, Miriam Llorian, Rocco D’Antuono, Anna Pavlova, Anna Maria Mavrigiannaki, Duncan McKenzie, Bethania García-Cassani, Maria Luisa Iannitto, Anett Jandke, Dmitry S. Ushakov, Adrian C Hayday

## Abstract

Because Tissue-Resident Memory T (T_RM_) cells contribute critically to body-surface immunoprotection and/or immunopathology in multiple settings, their regulation is biologically and clinically important. Interestingly, T_RM_ commonly develop in epithelia part-shaped by innate-like lymphocytes that become tissue-intrinsic during development. Here we show that polyclonal T_RM_ cells induced by allergic contact dermatitis (ACD) interact with signature intraepidermal γδ T cells, facilitating a feedback-loop wherein T_RM_-derived IFNγ upregulates PD-L1 on γδ cells that can thereupon regulate PD1^+^ T_RM_. Thus, T_RM_ induced by ACD in mice lacking either local γδ cells, or lacking a single gene (IFNγR) expressed by local γδ cells, displayed enhanced proliferative and effector potentials. Those phenotypes were associated with strikingly limited motility, reduced T_RM_ quality. and an impaired capacity to restrain melanoma. Thus, inter-individual and tissue-specific variation in how tissue-intrinsic lymphocytes integrate with T_RM_ may sit upstream of variation in responses to cancer, allergens and other challenges, and may likewise underpin inflammatory pathologies repeatedly observed in γδ-deficient animals.

## Introduction

Ongoing debates as to whether COVID-19 susceptibility is influenced by pre-existing memory T cells primed by common-cold coronaviruses simply highlights the critical importance of understanding the breadth of factors regulating the induction, turnover and quality of memory T cells (Bacher et al., 2020). However, despite extensive study, such factors are incompletely elucidated (Jameson and Masopust, 2018). The increasing acknowledgement that important memory responses are launched by Tissue-Resident Memory T (T_RM_) cells in peripheral extralymphoid sites such as the lungs, gut, or skin, highlights spatial constraints on memory T cell expansion and the need for memory cell turnover (Masopust and Soerens, 2019; Schenkel and Masopust, 2014). Clearly, with space being limited for T_RM_ cells of any one specificity, it is likely that mechanisms exist which promote the retention of cells of the highest quality in terms of affinity, durability, and rapid responsiveness to a recurrent challenge.

Local factors seem central to T_RM_ cell regulation (Hirai et al., 2021; Szabo et al., 2019; Takamura, 2018). Indeed, several datasets support a model by which T_RM_ progenitors first diverge from systemic T cells by acquiring generic, tissue-associated traits displayed by most T_RM_ compartments, after which they acquire site-specific traits, e.g. those distinguishing skin T_RM_ from lung T_RM._ (Bergsbaken and Bevan, 2015; Mackay et al., 2013; Schenkel and Masopust, 2014). Site-specific regulation would seem important given that this could equip T_RM_ cells to deliver functions appropriate to local anatomy (Takamura, 2018). Given that those anatomies, e.g. that of the epidermis, may show species-to-species variation, detailed aspects of site-specific regulation may not be perfectly conserved, but the underlying principles most likely will be.

Illustrating site-specific effects is epidermal T_RM_ regulation by keratinocytes (KC) (Hirai et al., 2019). Additionally, T_RM_ progenitors most often enter tissues structurally and functionally composed by heterogeneous cellular contributions. Thus, in addition to KC, T_RM_ maturing in the murine epidermis encounter tissue-intrinsic Langerhans cells (LC) and canonical Vγ5Vδ1^+^ TCRγδ^+^ dendritic epidermal T cells (DETC), each composing large, highly organised intraepithelial compartments that develop exclusively in the fetus, that display lifelong self-renewal, and that contribute to epidermal integrity (Chorro et al., 2009; Ikuta et al., 1990; Lewis et al., 2006; Park et al., 2021). Indeed, when viewed several weeks after a new T_RM_ compartment was established, DETC were observed to be spatially displaced by clusters of relatively motile CD8^+^ T cells (Zaid et al., 2014). However, the potential for these cells to meaningfully interact during the establishment of T_RM_ has not been examined.

In fact, a restraining regulatory role for γδ T cells is plausible given multiple long-standing but unexplained observations. First, DETC limit cutaneous graft-versus-host disease (GVHD) induced by inoculation of autoreactive CD4^+^ αβ T cells, and thereafter exclude subsequent CD4^+^ T cell inocula from becoming established in the same epidermal site, rendering the mice GVHD-resistant (Shiohara et al., 1996). Second, TCRγδ-deficient mice display exaggerated αβ T cell-dependent allergic contact dermatitis (ACD) responses, that could be completely prevented by reconstitution with fetal DETC progenitors (Girardi et al., 2002). Thus, in these qualitatively distinct settings, local γδ T cells limited the pathogenic potential of infiltrating αβ T cells. In this regard, inflammatory pathologies have been repeatedly observed in γδ T cell deficient mice, affecting several tissues from the gut to the testicles (Chen et al., 2002; Hayday and Tigelaar, 2003; Mukasa et al., 1995). Thus, one unexplored possibility is that T_RM_ may be regulated locally by tissue-intrinsic lymphocyte networks that in several sites comprise predominantly γδ T cells, but which in other sites might comprise innate-like αβ T cells, αβ T_reg_ cells, B1 B cells, and/or innate-lymphoid cells (ILC).

The murine epidermis is an attractive tissue in which to investigate local immune-mediated T_RM_ regulation, given that it is vitally important for barrier maintenance, is experimentally tractable, and is dominated at steady-state by only one tissue-intrinsic lymphocyte compartment: DETC. By focussing on ACD, a prevalent T cell-dependent pathology of humans that is commonly modelled in mice (Vocanson et al., 2006)(Girardi et al., 2002), we now show that several parameters of the murine polyclonal T_RM_ compartment are profoundly influenced by local γδ T cells. As one component of this crosstalk, T_RM_-derived IFNγ induced PD-L1 upregulation by intraepidermal γδ T cells: thus, akin to checkpoint blockade, removing DETC-mediated regulation slowed the motility of T_RM_, enhancing their potential to engage target cells and de-repressing their proliferative and effector potentials. However, the resultant T_RM_ phenotype was atypical, reflected in altered gene expression, and altered cell surface traits including reduced expression of CD5 that is a sentinel of reduced TCR affinity for antigen (Fulton et al., 2015; Mandl et al., 2013; Persaud et al., 2014). Associated with the altered quality of their T_RM_, γδ T cell deficient mice showed increased susceptibility to an epicutaneous melanoma ordinarily controlled by T_RM_ cells (Park et al., 2019). Thus, the local ecosystem into which T_RM_ cells enter includes heterotypic regulation by neighbouring tissue-intrinsic lymphocytes that may contribute to inter-individual, site-specific, and cross-species variation in immune responses to tissue-associated infections, inflammation, allergy, and cancer.

## Results

### ACD induces a polyclonal epidermal CD5^+^CD103^+^T_RM_ compartment

Mimicking frequent human exposure to chemical irritants, we employed a well-established system of allergic contact dermatitis (ACD) (Mackay et al., 2012) in which mice were initially sensitised to dinitrofluorobenzene (DNFB) on their abdomen, followed by re-exposure (“challenge”) ∼6 days later on their back or ear skin (Fig 1a, top panel). During sensitization, naive T cells in the draining lymph nodes (dLN) undergo antigen-driven clonal expansion and differentiate into effector cells, small numbers of which comprise systemic T_RM_ precursors. During challenge, those precursors accumulate at sites of allergen re-application, initiating a proinflammatory cascade, and developing into mature T_RM_ cells under the influence of local cues (Kok et al., 2020).

**Figure 1.**
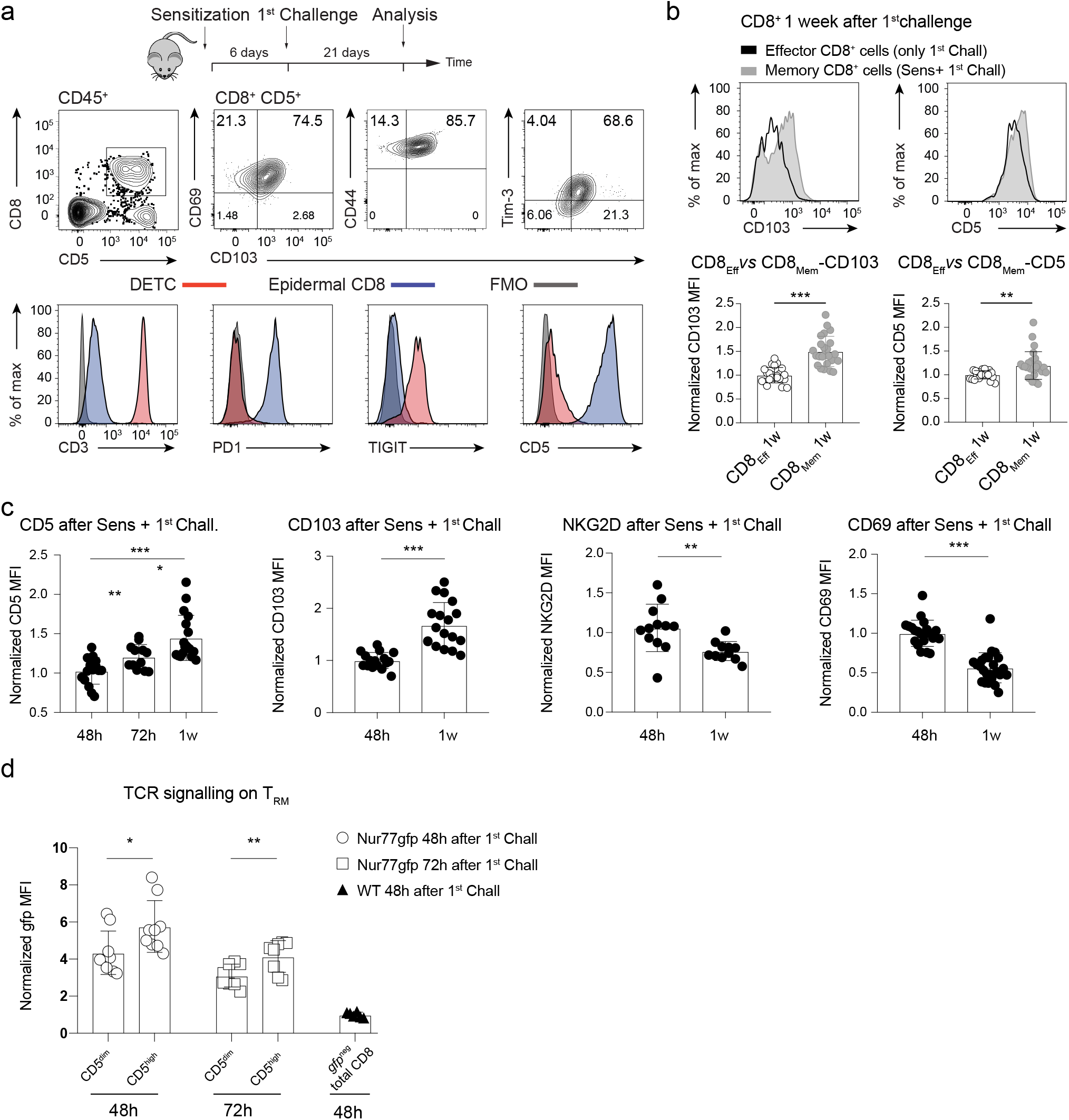
ACD induces a polyclonal epidermal CD5^+^CD103^+^T_RM_ compartment. **a)** ACD experimental protocol in WT mice (upper panel); flow cytometry evaluation of CD8^+^CD5^+^ T_RM_ for CD103, CD69, CD44 and Tim-3 (middle panel); flow cytometry analysis of epidermal CD103^+^CD8^+^ αβ T cells and DETC (lower panel). **b)** CD103 and CD5 expression assessed by flow cytometry on epidermal CD8^+^ T_RM_ cells and CD8^+^ T_Effector_ 1 week after either sensitisation and 1^st^ Challenge or primary antigen exposure (upper panel). Bar graphs show normalized MFI for CD103 and CD5 on CD8 T_Effector_ and CD8^+^ T_RM_ cells 1 week after DNFB treatment on ears (lower panel) (n = 24). **c)** Bar graphs show CD5, CD103, NKG2D and CD69 expression on epidermal CD8^+^ T cells after sensitisation and 1^st^ Challenge in WT mice at different time points. (n = 24). **d)** gfp levels on CD8^+^CD5^high^ *vs* CD8^+^CD5^low^ T cells, 48h and 72h after 1^st^ Challenge of Nur77^gfp^ mice (n = 8); “*gfp*^neg^ total CD8” refers to signal from non-transgenic WT mice. Data pooled from 3 biologically independent experiments. Data are mean□±□SD. Statistical analysis was performed using Student’s t-test. ***p < 0.001, **p < 0.01, *p < 0.05.

When we examined mice 21 days post-challenge (Fig 1a), epidermal T cells primarily comprised three subsets: a large population of tissue-intrinsic CD5^−^CD8^−^ DETC (Fig 1a, middle panels, leftmost plot); a relatively minor population of epidermal CD5^hi^ CD4^+^ cells (that was not studied further); and a largely homogeneous epidermal CD8^+^ population that was mostly CD103(αEβ7)^+^CD69^+^CD44^+^TIM3^lo^ (Fig 1a, middle panels), fully consistent with reported T_RM_ signatures (Clarke et al., 2019; Park et al., 2018). The CD8^+^ T cells mostly displayed very low CD3 expression (possibly reflecting a conformational change, since TCRβ staining was not atypically low), relatively high PD1, low TIGIT (“T cell immunoreceptor with Ig and ITIM domains”), and broad but generally high levels of CD5, whereas DETC displayed essentially the opposite phenotype: CD3^hi^,PD1^lo^,TIGIT^hi^,CD5^−/lo^ (Fig 1a bottom panel). Hence, although the local environment is a major driver of T_RM_ differentiation, it does not impose a singular phenotype on all T cell subsets that it houses.

Further illustrating this point, epidermal CD8^+^ αβ T cells harvested 1 week after DNFB treatment of mice that had not been previously sensitised were more akin to T-effector cells than to T_RM_, displaying significantly less CD103 upregulation, and also lower CD5 expression (Fig 1b). Indeed, in sensitised and challenged mice, the transition from the effector phenotype to one of upregulated CD103 and CD5 and reduced NKG2D and CD69 expression occurred incrementally between 48h and 1-week post-challenge (Fig 1c).

Additional evidence that DNFB sensitization initiated the development of T_RM_ cells was provided by assessing circulating cells from DNFB-primed mice for their potential to respond to epithelium-associated cytokines, TGFβ and IL15, by eliciting cells with a tissue memory precursor phenotype. Indeed, when splenic CD8^+^ T cells were harvested 5 days’ post DNFB sensitization and exposed for 2 days to TGFβ and IL15 alone, a significantly greater fraction acquired CD103 expression (∼45%) relative to CD8^+^ T cells from unchallenged WT mice (∼15%)(Supp Fig 1a). Such maturation absolutely depended upon cytokine provision (Supp Fig 1b), which also promoted CD5 upregulation (Supp Fig 1b) mostly by CD103^+^ cells (Supp Fig 1c), and that was further enhanced by TCR stimulation (Supp Fig 1d). Of note, upregulation did not occur on KLRG1^+^ CD8^+^ T cells, consistent with evidence that the KLRG1^−^ pool (Herndler-Brandstetter et al., 2018) is the exclusive source of T_RM_ progenitors (Supp Fig 1e) (Kok et al., 2020).

Because higher CD5 expression reportedly marks T_RM_ with higher affinity TCRs (Fiege et al., 2019), we investigated its potential significance using *Nur77*gfp mice in which a GFP reporter is expressed from the TCR-regulated promoter of the *Nur77* gene, ablation of which impairs T_RM_ formation (Boddupalli et al., 2016). Thus, hemizygous *Nur77*gfp mice were DNFB-sensitised, challenged 6 days later, and epidermal CD8^+^ T cells examined 48h and 72h post-challenge. At both time points, GFP levels were significantly higher (up to 50% greater) in CD5^hi^ *versus* CD5^dim^ CD8^+^ T cells (Fig 1d), indicating that they had experienced stronger TCR-dependent signal transduction. Likewise RANK-ligand, another TCR-responsive gene (Wang et al., 2002), was expressed significantly more strongly (∼3-fold) by epidermal CD5^hi^CD8^+^ T cells *versus* CD5^dim^CD8^+^ T cells. Thus, we consider CD5 expression as a quantitative index of T cell quality as it relates to TCR responsiveness, and thereby a marker of the functional polyclonality of the induced T_RM_. Conversely, CD5^dim^CD8^+^ and CD5^hi^CD8^+^ cells expressed comparable levels of CD8 and CD103, that do not reflect TCR signaling (Supp Fig 1f).

To examine the local functional consequences of T_RM_ induction, we induced ACD in hemizygous YFP-Yeti (enhanced transcript for IFNγ) mice, in which YFP reports transcriptional activity from the IFNγ-encoding gene commonly expressed by CD8^+^ T_RM_. YFP staining was spontaneously evident in over one third of epidermal CD8^+^ T cells 96h post-challenge, whereas none was detectable among DETC, LC, or low numbers of tissue-intrinsic αβ T cells that can be distinguished from T_RM_ by their high expression of CD3 (Supp Fig 1g). In sum, DNFB sensitisation induced CD8^+^ progenitor cells that upon epicutaneous challenge incrementally matured into polyclonal, epidermal T cells that collectively displayed a signature T_RM_ phenotype, that were locally unique in their expression of IFNγ transcripts, and that displayed a gradation of CD5 expression consistent with a spectrum of TCR signal strengths (Fulton et al., 2015; Mandl et al., 2013).

### ACD induces a local **γδ** T cell response

We next investigated whether the induction of the T_RM_ state that we have characterised above was set against a backdrop of concurrent DETC activation. At steady-state, canonical DETC are held in an activated-yet-resting state by Vγ5Vδ1^+^ DETC-specific selecting elements Skint1 and Skint2, expressed by suprabasal KC (D.McKenzie and A.C.H.; *in the press*). However, RNAs encoding those proteins, particularly *Skint2*, were rapidly down-regulated in DNFB-treated mice (Fig 2a). Indeed, within 72h post-challenge in sensitized mice, at a time of heightened maturation of T_RM_ cells, as exemplified by CD5 and CD103 upregulation (above), most DETC displayed a fully activated phenotype reflected by increased sphericity and CD69 upregulation (Fig 2b, arrows; Supp Fig 2a, b). Moreover, this phenotype was exclusively that of epidermal γδ T cells in DNFB-challenged mice since those cells uniquely express Vγ5 and there was no evidence of epidermal infiltration by γδ T cells using other TCRs, e.g., Vγ4, which is common among dermal γδ T cells (Supp Fig 2c; (Jiang et al., 2017)).

**Figure 2.**
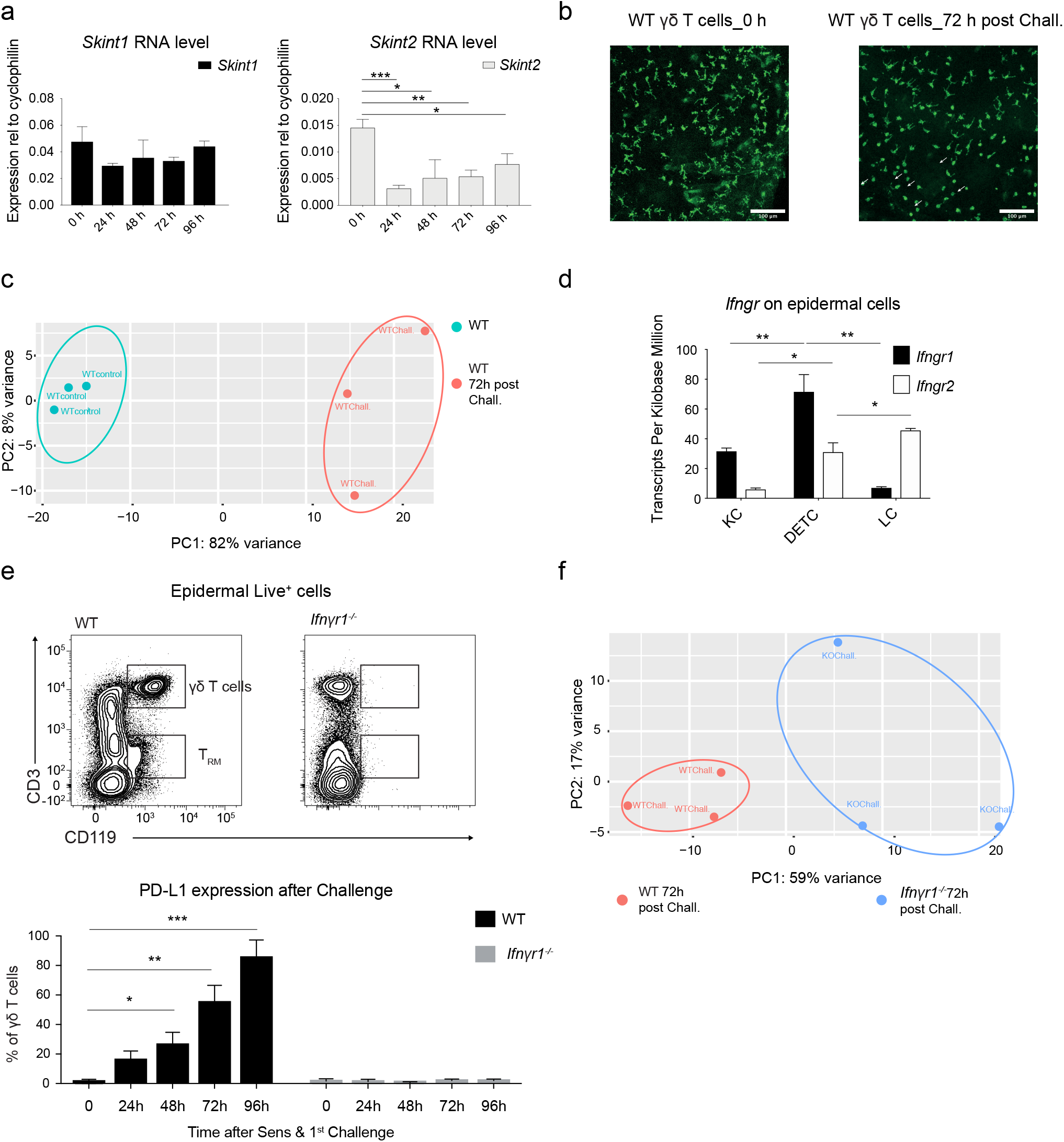
ACD induces a local γδ T cell response. **a)** *Skint1* and *Skint2* mRNA analysis by qPCR in back epidermis of adult mice normalised to Cyclophilin at different time points after sensitisation and 1^st^ challenge; h=hours, n = 3. Data are mean□±□SD of a representative experiment of two independent experiments. **b)** Comparison of ear skin DETC stained for TCRγδ (green) and assessed by confocal microscopy in WT mice either unchallenged or 72h after 1^st^ challenge. Scale bar 100 µm. Representative of 3 independent experiments. **c)** PCA of bulk RNAseq analysis of epidermal γδ T cells sorted from unchallenged or challenged (72h after 1^st^ challenge [“Cha”]) mice; colour denotes status (n=3 n’=3). **d)** Epidermal cells from unchallenged mice were sorted and *Ifngr1* and *Ifngr2* gene expression determined from RNAseq on the indicated cell types (n=3). **e)** Representative flow cytometry of CD3 and CD119 (IFNγR1) expression by live epidermal cells from WT and *Ifn*γ*r1^−/−^*animals (3 independent experiments) (upper panel). Frequency of epidermal γδ T cells expressing PD-L1 as assessed by flow cytometry at different time points after sensitisation and 1^st^ Challenge on WT and *Ifn*γ*r1^−/−^*mice (bottom panel). **f)** PCA of bulk RNAseq analysis of epidermal γδ T cells sorted 72h after Sensitisation and 1^st^ Challenge of either WT or *Ifn*γ*r1^−/−^* mice; colour denotes status (n=3 n’=3). **a-d-e)** Statistical analysis was performed using one-way ANOVA with Sidak’s multiple comparisons post-hoc test. ***p < 0.001, **p < 0.01, *p < 0.05.

At a higher level of resolution, there were profound differences in the gene expression profiles of Vγ5Vδ1^+^ DETC recovered 72h post-challenge of sensitised mice *versus* those at steady-state, as illustrated by a Principal Component Analysis (PCA) of an RNAseq analysis (Fig 2c; Supp Table 1; GSE164023). Thus, by 72h many mRNAs associated with T cell activation were increased, including those encoding cytokines, cytokine receptors, and selected chemokines; those encoding costimulatory/coinhibitory receptors, such as 4-1BB, Icos, and Tim3; and those encoding cytolytic mediators (Supp Fig 2d). Also, conspicuously upregulated were IFNγ-responsive genes, including *Ly6a, Stat1, Cxcl10* (encodes IP10), and *Cd274* (encodes PD-L1) (Supp Fig 2d). Strikingly, by comparison to KC and LC, DETC were the epidermal cells that most consistently expressed RNA and protein for both chains of the IFNγR that are each required for IFNγ-responsiveness (Krause et al., 2002) (Fig 2d; GSE160477; Supp Fig 2e). Consistent with this, the DETC response to IFNγ, measured by *Sca1* upregulation (Ma et al., 2001) was significantly greater than the LC response (Supp Fig 2f).

In this context, DETC uregulation of PD-L1, which was incremental over 96h following ACD induction, did not occur in *Ifnγr1*^−/−^ mice (Fig 2e, upper and lower panels). Indeed, DETC sampled at 72h DNFB post-challenge from WT mice showed substantially different transcriptomes *versus* those sample from *Ifnγr1^−/−^* mice, as revealed by PCA (Fig 2f; Supp Table 2; GSE164023). For example, activated *Ifnγ*r1-deficient DETC showied reduced expression of *Stat1, Tbx21* (encodes Tbet), and *Ly6a*, as well as ablated PD-L1 (Supp Table 2). These data establish that at early time points during ACD induction *in vivo*, IFNγ is a major, heretofore under-appreciated regulator of DETC in an environment in which IFNγ RNA is primarily expressed by T_RM_ (above) and DETC are the strongest expressers of IFNγ receptors.

### DETC-T_RM_ Interactions *in vivo*

Based on the data obtained, we hypothesised that rather than inhabiting parallel universes, locally developing T_RM_ and activated tissue-intrinsic γδ T cells might be connected by an IFNγ-dependent feedback loop. This would be consistent with a “sensing and alarm” function by which T_RM_ condition the immunological tenor of local tissue (Schenkel et al., 2013). Investigating this further, we observed that when whole epidermal preparations from ACD-challenged mice were harvested and rested for 48h, ≥80% of TCRγδ^+^ cells expressed PD-L1, consistent with the data described above, whereas when γδ T cells were maintained in a parallel culture following flow cytometry-based purification, most became PD-L1^−^ (Supp Fig 3a). Thus, we considered that PD-L1 expression might reflect sustained exposure to IFNγ *in vivo.* Moreover, and by contrast to many freely diffusible cytokines, IFNγ is reportedly most active within the direct proximity of its cellular source (Krummel et al., 2018). Based on these considerations, we hypothesised that direct interactions might commonly occur between T_RM_ and DETC. Although very few T_RM_ and DETC interactions were observed in the epidermis following *Herpes* virus-induced T_RM_ formation (Zaid et al., 2014), those analyses were undertaken at steady-state, long after the establishment phase of a T_RM_ compartment which is the time-frame considered here.

To detect any direct DETC-T_RM_ interactions, we applied confocal microscopy to Yeti mice at 72h post-challenge, detecting YFP as a surrogate of IFNγ-producing cells, and staining with antibodies against CD8 and TCRγδ, respectively. High densities of CD8^+^ T cells were visualised at the challenge site, many of which expressed YFP (that could be sufficiently bright as to diminish the CD8 signal) (Fig 3a, red). Conspicuously, many of these cells were in intimate juxtaposition with DETC (Fig 3a, blue; arrowed; Fig 3b, white; Supp Fig 3b; Supp Video1). Furthermore, DETC that were detected interacting with CD8^+^ T cells also expressed PD-L1 (Fig 3c).

**Figure 3.**
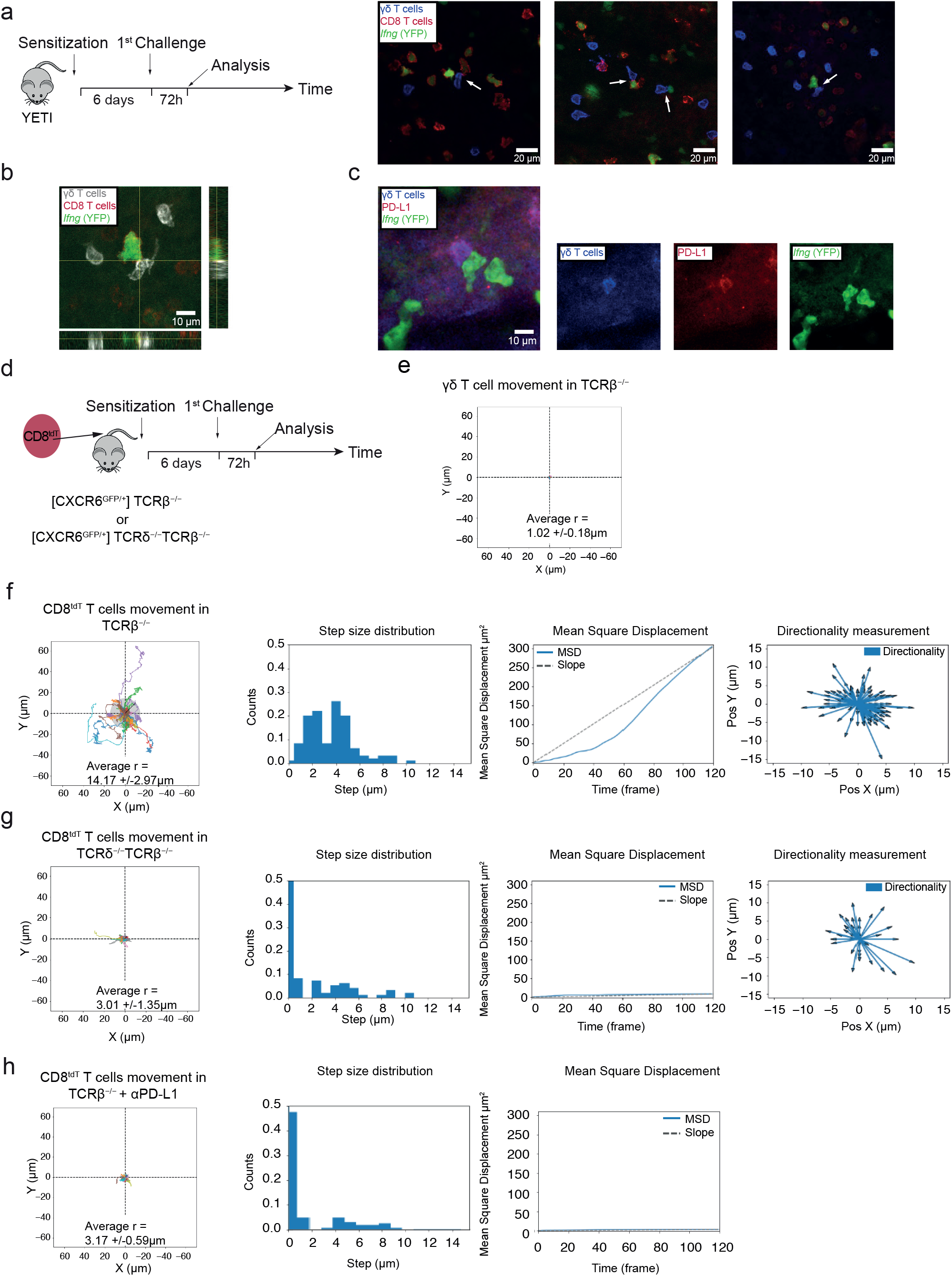
DETC-T_RM_ Interactions *in vivo*. **a-b)** ACD experimental protocol (left panel). Confocal microscopy images of ear epidermal sheets of YFP-Yeti mice 72h after Sensitisation and 1^st^ Challenge. DETC stained for γδ TCR (blue or grey) and CD8 T cells for CD8α (red). Scale bar as indicated in each image□(µm) (n=3). **c)** Confocal microscopy images of ear epidermal sheets of YFP-Yeti mice 72h after Sensitisation and 1^st^ Challenge. DETC stained for γδ TCR (blue or grey) and PD-L1 (red). Scale bar as indicated in each image (µm). **d)** Experimental protocol for tdTomato-CD8^+^ adoptive transfer to [CXCR6^GFP/+^]TCRβ^−/−^ or [CXCR6^GFP/+^]TCRβ^−/−^TCRδ^−/−^ mice, followed by ACD induction**. e)** Confocal Microscopy cell migration tracks of individual CXCR6^GFP/+^ γδ T cells in the skin 72h after sensitisation and 1^st^ challenge normalized for their origin. **f-h)** Confocal Microscopy cell migration tracks of individual tdTomato T_RM_ in the skin 72h after sensitisation and 1^st^ challenge normalized for their origin. The grey circle radius is the weighted mean of the track total displacements. Reported values for “r”: weighted mean and SD of the weighted mean. **f)** [CXCR6^GFP/+^]TCRβ^−/−^ mice: Total of 26 cells in 3 different experiments. **g)** [CXCR6^GFP/+^]TCRβ^−/−^TCRδ^−/−^ mice: Total of 22 cell in 3 different experiments. **h)** [CXCR6^GFP/+^]TCRβ^−/−^ mice + aPD-L1: Total of 20 cells in 3 different experiments. **f-g-h)** T_RM_ movement analysis for tdTomato-CD8^+^ adoptively transfer to [CXCR6^GFP/+^]TCRβ^−/−^, [CXCR6^GFP/+^]TCRβ^−/−^TCRδ^−/−^ mice or [CXCR6^GFP/+^]TCRβ^−/−^ mice + aPD-L1: Step size distribution, Mean Square Displacement and directionality quantified using Manual Tracking plugin in FIJI (Schindelin et al., 2012) (lower panels). One representative experiment of the 3 experiments performed is shown.

A potent, relatively underexplored consequence of PD-L1/PD1 regulation is an increased motility of PD1^+^ T cells. In the context of Type I diabetes, enhanced PD-L1-dependent motility of pancreatic islet antigen-specific TCRαβ^+^ cells within tissue-draining lymph nodes was interpreted as promoting tolerance by restricting antigen/TCRαβ-dependent, “strong-stop” contacts with dendritic cells to only cells with the highest affinity TCRs; i.e. the highest quality, which seldom defines autoreactive T cells (Fife et al., 2009). Likewise, in the context of oncogene-induced tissue injury, enhanced PD1-dependent motility could beinterpreted as promoting tolerance by limiting αβ T cell interactions with damaged and proliferating cells during tissue regeneration (Kortlever et al., 2017). We therefore examined whether epidermal CD8^+^ T cells showed a signature pattern of motility under the influence of DETC and PD-L1.

To this end, we employed CXCR6-gfp mice in which DETC at steady-state are GFP^+^ as are tissue-infiltrating CD5^+^CD8^+^ αβ T_RM_ following challenge (Supp Fig 3c). Those mice were crossed to homozygosity for TCRβ-deficiency, so that DETC were the only GFP^+^ cell type in the skin, and those mice were reconstituted with CD8^+^ αβ T cells derived from TdTomato mice so that they too could be visualized in recipients(Fig 3d). 72h after DNFB-sensitized mice were challenged with DNFB, ears were assessed by confocal microscopy. The green cells (DETC) were essentially sessile, as previously reported (Park et al., 2021), with the motility of tracked cells being captured over a time-frame of 120 seconds by a circle with an average radius of 1.02 +/− 0.18μm (Fig 3e). By contrast, red (CD8^+^) cells were motile, captured in the same time-frame by a circle of radius 14.17 +/− 2.97μm (Fig 3f). Moreover, there were clear instances of motile CD8^+^ T cells making episodic and repeated interactions with DETC (Supp Video 1). Analysed in greater detail, the CD8^+^ cells showed an average stepsize (the distance measured in each 30 sec interval) of ∼3.5um and no skewed directionality. Nonetheless, their mean square deplacement (MSD) from their start positions did not increase linearly with time, with some restraint on motility evident particularly in the early imaging intervals by the negative deviation from the straight line (Fig 3f; Supp Video 2).

To better understand this motility pattern, we set up simulations in which a red motile cell was positioned among sessile green cells distributed to mimic the positions of those real events observed in the tissue. We assigned value functions reflecting the random movement of the cells in terms of displacement (0.0-1.0) and direction (0°- 360°); their hypothetical activation by an initiation factor, which would reflect the response to PD-L1 engagement and which would increase stepsize (0.99); and their hypothetical attraction to a nearest neighboring DETC within 100μm (0.7) that would affect cell direction. Additionally, we imposed a maximum stepsize of 14μm based on our real events observations. Additionally, the model included terms to simulate potential contacts between the two cell populations (defined as where two cells are less than 3μm away) that might account for reduced stepsize and hence the displacement from linear MSD. This simulation (Supp Fig 3d) strikingly phenocopied the data obtained from tracking epidermal CD8^+^ T cells following DNFB challenge, strongly attesting to the validity of assigning activation and attraction functions for CD8^+^ T cells within 100μm of DETC. Moreover, by observing 40 CD8^+^ T cells in a given field-of-view containing sessile DETC, the model permitted us to estimate that ∼25% of CD8^+^ T cells would be in contact with DETC over the 120 sec time-frame (Supp Fig 3e).

We therefore hypothesized that in the absence of γδ T cells, CD8^+^ T cell movement would be significantly altered by a lack of a PD-L1-dependent activation function, and a lack of an attraction function. To test this hypothesis, we repeated the study by using TdTomato CD8^+^ αβ T cells to reconstitute CXCR6.gfp TCRβxδ^−/-^ mice that lacked all γδ T cells, and that were then sensitized and challenged. Indeed, at 72h post challenge, the CD8^+^ T cell motility (r = 3.01 +/− 1.35mm) (Fig 3g) was much reduced relative to that observed in γδ-suffcient mice (see Fig 3f). Furthermore, the modal stepsize was zero, as a result of which there was negligible MSD (Fig 3g), with the rare instances of motility showing no directional skewing.

When we then simulated a situation in which the activation and attraction functions were both negligible, the outcome strikingly phenocopied that observed for CD8^+^ T cell motility in the absence of γδ T cells (Supp Fig 3f). Having established the dependence of T_RM_ motility on γδ T cells. We next asked whether it was PD-L1-dependent, by administering anti-PD-L1 antibody 24h prior to challenge of CXCR6.gfp TCRβ^−/-^ mice reconstituted with TdTomato CD8^+^ T cells. Indeed, the cells’ movement was greatly impaired by anti-PD-L1 (r =3.17 +/− 0.59μm; modal stepsize = 0) (Fig 3h), closely phenocopying the impact of γδ T cell deficiency. Interestingly, this treatment also phenocopied the impact of γδ T cell deficiency (Girardi et al., 2002) in leading to increased ear swelling responses to DNFB-challenge (Supp Fig 3g), as was also reported in *Pdl1^−/-^* mice (Hirano, 2021).

### **γδ** T cell-mediated regulation of the T_RM_ phenotype

As considered above, DNFB-stimulated ACD applied to γδ T cell-deficient mice provoked highly exaggerated αβ T cell-dependent inflammation (Girardi et al., 2002), that we now hypothesise reflects an impact on T_RM_ cells. We therefore made a more detailed examination of the impact of epidermal γδ T cells on T_RM_ development *in situ.* To this end, we compared the phenotypes of T_RM_ cells at 72h after a 2^nd^ DNFB challenge of WT mice *versus* age-matched Tac mice which specifically lack canonical Vγ5Vδ1^+^ DETC because of a mutation in *Skint1*, the obligate DETC-selecting determinant (Barbee et al., 2011; Lewis et al., 2006). Note that Tac epidermis is replete with “replacement DETC” comprising non-Vγ5Vδ1^+^ γδ T cells, so that T_RM_ in both strains develop amidst tissue-intrinsic γδ T cells. Nonetheless, the replacement non-Vγ5Vδ1^+^ γδ T cells were not substantially activated during ACD, by comparison to canonical Vγ5Vδ1^+^ γδ T cells, as illustrated by their limited CD69 response (Supp Fig 4a).

Based on the phenotypic analysis of T_RM_ presented above, freshly isolated CD8^+^ T cells were flow-sorted for CD45^+^TCRβ^+^CD103^+^CD5^+^ and subjected to 3’ mRNA single-cell transcriptomics, obtaining 483 mean reads per cell for the WT sample and 1346 for the Tac sample, with the top 1000 most variable genes selected for each. Sequencing saturation was >90% for all samples, indicating comprehensive sampling of available transcripts. After pre-processing, normalization and batch correction, integrated analyses were applied to discriminate common cell types and facilitate comparative analyses, as was described (Stuart et al., 2019). The application of t-distributed stochastic neighbour embedding (t-SNE) to a mixed data-set from challenged WT and Tac mice permitted five distinct lymphocyte clusters to be discriminated according to their aggregate gene expression profiles (Fig. 4a).

**Figure 4.**
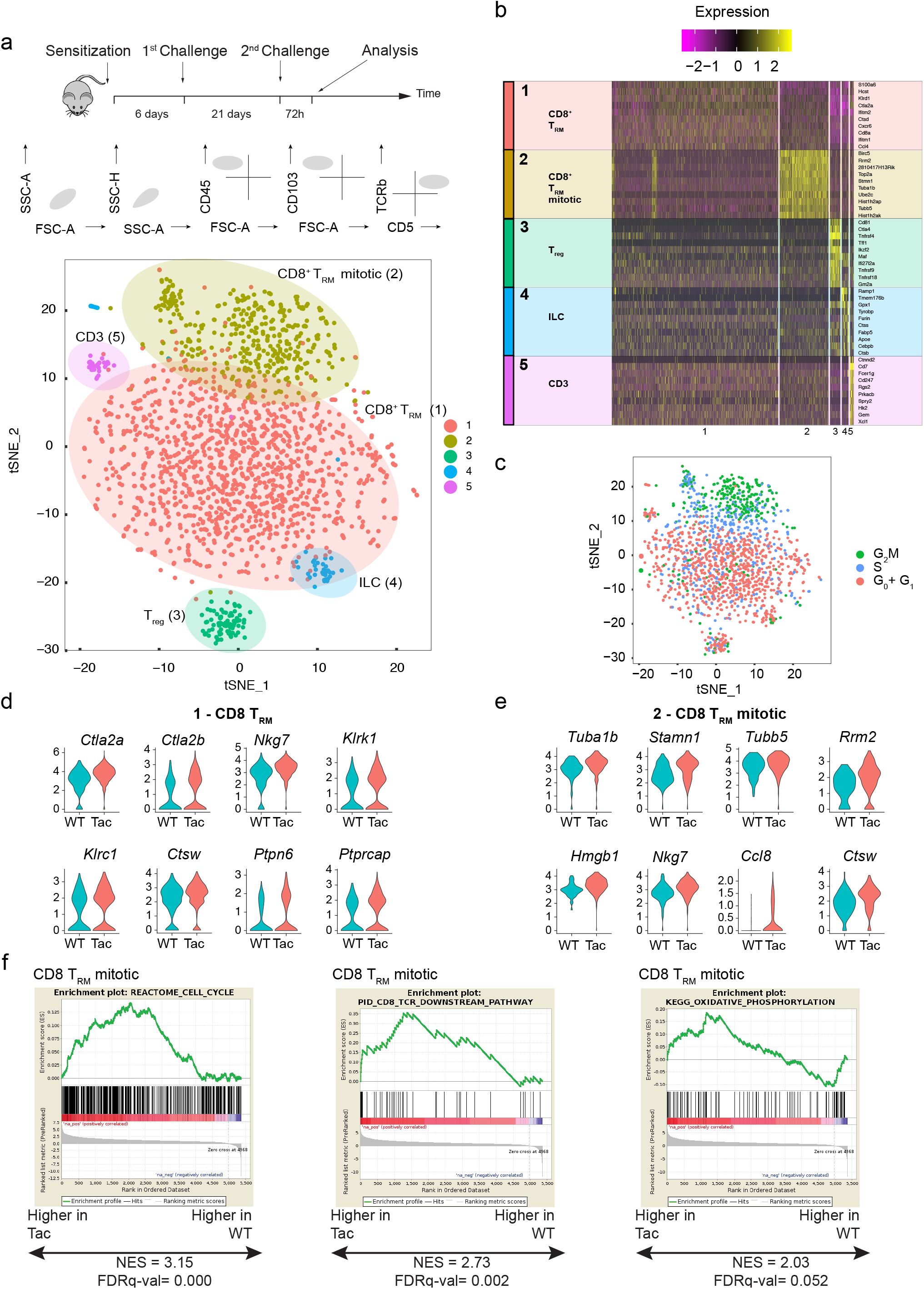
γδ T cell-mediated regulation of the T_RM_ phenotype. **a-f)** scRNAseq analysis of CD8 T cells isolated from WT and Tac mice 72h after 1^st^ challenge. **a)** ACD experimental protocol and workflow for isolating T cells (upper). t-SNE plot with colours demarcating 5 distinct clusters based on gene expression differences for 1829 cells passing quality control (WT *+* Tac). The numbers in parentheses correspond to the clusters listed in b. **b)** Heat map with cells grouped into clusters (indicated by coloured bars at the left). The 10 genes most highly differentially expressed by each cluster are denoted on the right. **c)** t-SNE plot of scRNAseq dataset showing Seurat cell cycle analysis. **d)** Violin plots comparing the expression of genes associated with proliferation in single cells from WT (green-blue) versus Tac (pink) mice. **e)** Violin plots comparing the expression of genes associated with CD8^+^ T cell effector functions in single cells from WT (green-blue) versus Tac (pink) mice. **d**-**e)** Shown are violin-shaped fitting areas for statistically significant differences in gene expression of T_RM_ from WT *vs* Tac mice. ***p < 0.001. **f)** Gene set enrichment analysis (GSEA) of T_RM_ mitotic cluster from Tac *vs* WT backgrounds using indicated public gene sets. The enrichment score (NES) and p value are reported.

For the purposes of illustration, the ten genes whose expression was most enriched in the respective clusters are denoted in Fig. 4b, whereas the complete information is provided online (GEO accession number GSE164023). The most prominent clusters were two *Cd8α*^+^ T cell clusters, “Cluster 1” and “Cluster 2”, further considered (below); Cluster 3 comprised a small group of *Cd4*^+^ T cells enriched in T_reg_-associated transcripts; Cluster 4 comprised contaminating *Cd3*^−^*TCR*β^−^ ILC; and Cluster 5 comprised a very minor group of *Cd3-*expressing cells of uncertain classification (they were not TCRγδ^+^).

Accepting a false discovery rate (FDR) of □<1%, Cluster 1 cells composed the largest cluster (Fig. 4b: GSE164023). The cells expressed a prototypic T_RM_ signature conserved in mice and humans that included *Cxcr6*, *Cd8a, Itgae* (encoding CD103), *Pdcd1* (encoding PD1), *Ccl4* and *Cd69* (Fig 4b, Supp Fig 4b) (Mackay et al., 2013; Schenkel and Masopust, 2014). Additionally, genes associated with CD8^+^ T cell effector responses were highly expressed, including *Gzmb, Icos, Tnf* and *Ifng*. (Supp Fig 4b). Conversely, Cluster 1 showed scant expression levels of genes associated with lymph node-homing and T_EM_ cells, including *sell* (encoding CD62L), *ccr7, s1pr1* and *s1pr5*, *klrg1* and *klf2* (Fig 4b, Supp Fig 4c; GSE164023) (Kumar et al., 2017; Mackay et al., 2013; Schenkel and Masopust, 2014).

In many respects Cluster 2 cells shared the collective Cluster 1 T_RM_ signature (Fig 4b, Supp Fig 4a; GSE164023). However, as is clear from Fig 4b and Supp Fig 4b, Cluster 2 was uniquely enriched in RNAs associated with active cell cycling, including *Stmn1* (encoding stathmin1), *Tubb5*, *Rrm2* (encoding ribonucleoside-diphosphate reductase subunit M2), and *Mki67* (encoding Ki67 which is a marker of cycling cells) (Fig 4b, Supp Fig 4b) (Savas et al., 2018). Moreover, Cluster 2 was strongly enriched in RNAs associated with the S and particularly the G_2_-M phases of the cell cycle (Fig. 4c). Thus, the epidermis at 72h post 2^nd^ challenge concurrently housed proliferative (Cluster 2) and resting (Cluster 1) CD8^+^ T_RM_ cells.

We next compared the distribution across these clusters of epidermal cells recovered from Tac *versus* WT mice 72h post challenge (Supp Fig 4c, d). There was a relative loss in Tac of cells from minor Clusters 3 and 4 which evidently reflects fewer contaminating T_reg_ and ILC. Beyond this, the major Clusters 1 and 2 were densely populated in Tac mice and their relative ratios were comparable with those in WT. Nevertheless, Tac-derived T_RM_ showed significant quantitative shifts in the expression of specific gene-sets: for example, Tac-derived Cluster 1 cells showed increased expression of genes associated with cytolytic effector functions, including *Ctla2a, Nkg7, Ctsw, Klrk1*, and signalling status including *Ptpn6* (encodes Shp1) and *Ptprcap*, both of which encode regulators associated with modulation of T cell activation (Cho et al., 2016; Dong et al., 2006) (Fig 4d).

Likewise, Tac Cluster 2 T_RM_ also showed increases in genes associated with cytotoxicity including *Ctsw* and *Nkg7*, and in genes associated with outcomes of TCR signalling and oxidative phosphorylation (Fig 4e,f). Additionally, Tac Cluster 2 T_RM_ showed significant increases relative to WT Cluster 2 in mitosis-associated genes, including *Stmn1, Tuba1b, Tubb5, Rrm2,* and *Hmgb1*, whereas they showed relatively reduced expression of genes associated with responsiveness to cytokines and overall immune stimluation (Supp Fig 4e). In sum, the transcriptomic data indicated that in mice lacking canonical γδ DETC, polyclonal epidermal CD8^+^ T_RM_ cells had: first, greater cytolytic effector potentials, akin to CD5^dim^NKG2D^hi^ T-effector cells observed in mice exposed to DNFB without prior sensitisation (see Fig 1); and second, more proliferative activity, as has been associated with CD5^lo^ T_RM_, which commonly have lower TCR affinity relative to CD5^hi^ T_RM_ (Fiege et al., 2019; Voisinne et al., 2018).

### IFN**γ**-dependent T_RM_ regulation by local **γδ** T cells

To further examine the status of T_RM_ developing in γδ DETC deficient mice, the sensitisation and challenge protocol was applied to three strains: Tac, as used above; TCRδ^−/−^ that lacks all γδ T cells; and Vγ5Vδ1^−/−^ that cannot assemble the canonical DETC TCR. Like Tac mice, Vγ5Vδ1^−/−^ mice harbour “replacement” γδ T cells expressing a range of non–canonical γδ TCRs. Conversely, replacement DETC in TCRδ^−/−^ mice comprise TCRαβ^+^CD8^+^ cells that are clearly distinguishable from CD8^+^ T_RM_ because they are CD8^lo^ and do not express CD5 (Fig 5a; compare upper with lower panels). The percentage of epidermal T cells occupied by CD8^lo^CD5^+^ T_RM_ was often conspicuously greater in DETC-deficient mice and this was further exaggerated following a second DNFB challenge three weeks after the first (Fig 5a, b): indeed, absolute epidermal T_RM_ counts were significantly greater in TCRδ^−/−^, Vγ5Vδ1^−/−^ and Tac strains, albeit that their distributions overlapped with WT counts (Fig 5c). Note, because of replacement DETC, one cannot simply attribute greater T_RM_ expansion in γδ-deficient settings to there being an empty niche to fill; a possibility that is also excluded by observations described below.

**Figure 5.**
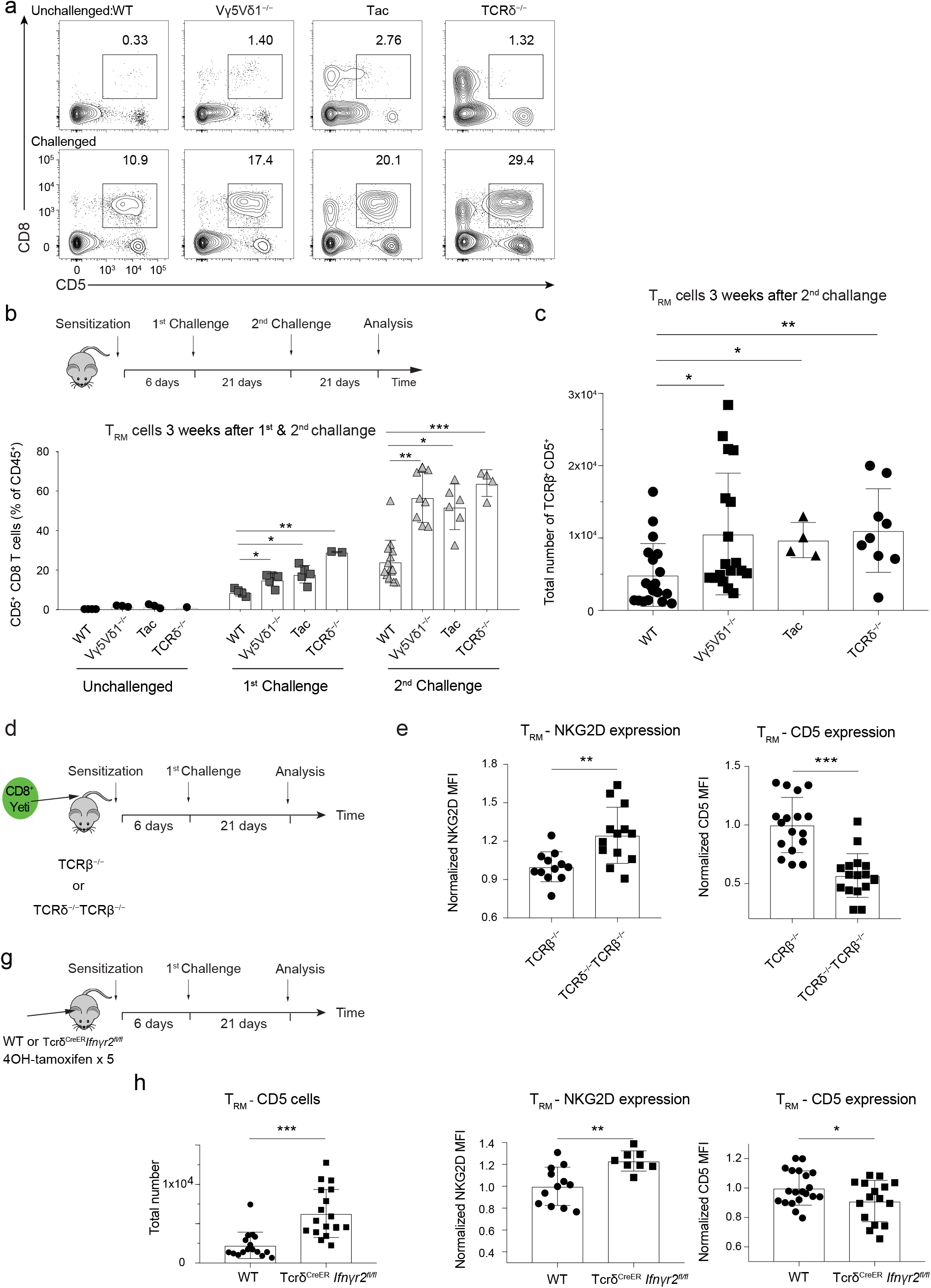
IFNγ-dependent T_RM_ regulation by local γδ T cells. **a)** Representative flow cytometry data of CD8 and CD5 expression by epidermal CD45^+^ cells from unchallenged (upper panels), or 21 days after sensitisation and 1^st^ challenge (lower panels) in WT, Vγ5Vδ1^−/−^, Tac and TCRδ^−/−^ mice. **b)** Frequency of T_RM_ from WT, Tac, Vγ5Vδ^−/−^ and TCRδ^−/−^ mice following indicated challenges. **c)** Total counts of TCRβ^+^CD5^+^ cells 21 days after the 2^nd^ Challenge in WT, Tac, Vγ5Vδ1^−/−^ and TCRδ^−/−^ mice. **b-c)** Data pooled from n = 3 biologically independent experiments with a total of 40 mice. Data are mean□±□SD. Statistical analysis was performed using one-way ANOVA with Sidak’s multiple comparisons post-hoc test. ***p < 0.001, **p < 0.01, *p < 0.05. **d)** ACD experimental protocol following adoptive transfer of CD8^+^ T cells from YFP-Yeti mice to TCRβ^−/−^ and TCRδ^−/−^TCRβ^−/−^ recipients. **e)** Bar graphs show normalized CD5 and NKG2D surface expression of transferred CD8^+^ cells from TCRβ^−/−^or TCRδ^−/−^TCRβ^−/−^ recipients at 21 days post-challenge. Data pooled from 3 biologically independent experiments with a total of 33 mice. **g)** Experimental protocol for deletion of *Ifn*γ*r2* gene specifically from γδ T cells, followed by ACD induction. **h)** Bar graphs show normalized T_RM_ surface expression of CD5 and NKG2D and T_RM_ quantification assessed by flow cytometry 21 days after 1^st^ Challenge in WT and *Tcrd^creER^Ifn*γ*r2^fl/fl^* mice. Data pooled from 3 biologically independent experiments with a total of 36 mice. **e-h)** Data are mean□±□SD. Statistical analysis was performed using Student’s t-test. ***p < 0.001, **p < 0.01, *p < 0.05.

To further investigate the formation of T_RM_ in γδ DETC-deficient mice, and to avoid the issue of replacement γδ T cells, we adoptively transferred CD8^+^ T cells from Yeti mice to TCRβ^−/−^ (above) or TCRδ^−/−^TCRβ^−/−^ mice that were then sensitised later and subsequently challenged, following the protocol described for the imaging studies, above (Fig 5d). Strikingly, epidermal CD8^+^ cells recovered from γδ T cell-deficient recipients were qualitatively distinct from those repopulating γδ T cell-sufficient mice, in that they expressed significantly higher levels of T-effector markers including NKG2D, NKG2A, ICOS, and 4-1BB and their YFP-fluorescence was higher indicating increased *Ifng* transcription (Fig 5e; Supp Fig 5a). Of note, these results were then validated in mice specifically lacking Vγ5Vδ1 DETC (Supp Fig 5b). Strikingly, T_RM_ maturing in γδ DETC-deficient mice also expressed significantly lower levels of CD5 (Fig 5e; Supp Fig 5c), as was the case for epidermal T-effector cells maturing in mice exposed once to DNFB without prior sensitisation (above). Pursuing this further, we observed that local inoculation of WT mice with the squamous cell carcinoma line, PDV (Girardi et al., 2004) provoked a major epidermal influx of CD5^+^ CD8^+^ T cells (Supp Fig 5d). When TCRδ^−/−^ mice were the recipients of PDV cells, the induced epidermal CD8^+^ T cells again expressed CD5 (clearly distinguishing them from replacement DETC), but they were mostly CD5^lo^ by comparison to the proportions of CD5^hi^ cells seen in WT recipients (Supp Fig 5e).

Hence, the importance of γδ T cells for driving the upregulation of CD5 on epidermal CD8^+^ T cells was evident in two entirely independent systems. We therefore asked whether such an affect could be observed *in vitro* during the cytokine-dependent maturation of T_RM_ from splenocyte progenitors, described above (see Supp Fig 1) Strikingly, co-culture with short-term lines of Vγ5Vδ1^+^ DETC significantly increased CD8^+^ T cell maturation to CD103^+^ and those cells expressed a significantly higher amount of CD5 (Supp Fig 5f). Interestingly, CD5^lo^ *versus* CD5^hi^ CD8^+^ T cells were reported to respond less well to cytokines and to show increases in outcomes associated with TCR signalling (Cho and Sprent, 2018), which were precisely those traits deduced from the transcriptomic analysis of T_RM_ in γδ-deficient mice (above).

We then asked whether these examples of T_RM_ regulation by local Vγ5Vδ1^+^ γδ T cells reflected at least in part the cells’ regulation within an IFNγ-driven loop, as hypothesised above. Indeed, *Ifnγr1^−/−^* mice phenocopied DETC-deficient mice in response to ACD, displaying a significantly greater frequency of T_RM_ post-2^nd^ challenge (Supp Fig 5g). Indeed, somewhat counter intuitively, IFNγR1-deficient mice are prone to inflammation akin to γδ-deficient mice (Willenborg et al., 1996, 1999). Building on these observations, we next generated a γδ-specific IFNγR2-deficient mouse strain by crossing a floxed allele of *Ifnγr2* with a mouse expressing a tamoxifen-responsive cre-recombinase (CreER) under the control of the TCRδ promoter (Fig 5g), so as to preclude PD-L1 upregulation specifically in DETC. As expected, DETC recovered from those mice following tamoxifen induction of *Ifnγr2* deletion (Supp Fig 5h) were greatly impaired in their capacity to upregulate *Sca1* in response to recombinant IFNγ (Supp Fig 5i).

Strikingly, the ACD response of these mice phenocopied the ACD response of DETC-deficient mice and *Ifnγr^−/−^* mice in that there was an exaggerated amplification of T_RM_ cells which expressed lower levels of CD5 and higher levels of effector T cell-associated proteins including NKG2D (encoded by *Klrk1*) (Fig 5h). Thus, the T_RM_ phenotype was significantly affected by the targeted deletion of a single gene in an heterologous cell type (DETC) with which maturing T_RM_ evidently interact (above). Of note, because canonical DETC are still present in *Tcrd^cre^Ifnγr2^fl/fl^* mice, these results further diminish any capacity to use spatial arguments to explain DETC-dependent limitation of T_RM_ expansion. Moreover, the data evoke other settings in which T_RM_ regulation by PD-L1 has been reported. For example, genetic disruption and intracerebroventricular blockade of PD-1/PD-L1 signalling increased the number of murine polyoma virus-specific CD8^+^ T_RM_ cells, the fraction of those cells expressing CD103, and the effector capacity of such cells (Shwetank et al., 2019), collectively mimicking our observations of the shared impacts of DETC deficiency and IFNγR deficiency on T_RM_ cells.

Finally, we cultured T_RM_ *in vitro* with very low concentrations of the activating anti-TCRβ antibody, H57, with and without short-term DETC cultures (above) and with and without anti-PD-L1, for which there was an isotype control. Those studies showed that the numbers of T_RM_ were significantly reduced by co-culture with DETC, but that this effect was overcome by anti-PD-L1 (Supp Fig 5j): findings consistent with the enrichment for more highly proliferative T_RM_ in settings of γδ-deficiency, IFNγR-deficiency, and PD-L1 blockade, above.

### γδ T cell-deficiency impairs T_RM_-regulated tumour control

The findings reported in this study collectively demonstrate that in response to DNFB-challenge of pre-sensitised mice, the phenotype of the polyclonal T_RM_ compartment that forms is heavily influended by local γδ T cels, which in particular promote a T_RM_ phenotype of increased CD5 expression that is associated with higher affinity cells, *versus* a proliferative effector phenotype more commonly seen after single episodes of antigen exposure. These different phenotypes are germane to ongoing debates as to whether T-effector or T_RM_ TCRαβ ^+^ cells are most germane to tumour control. To examine this, we employed with slight modification a system in which control over melanoma growth depends on epidermal T_RM_ cells (Park et al., 2019).

Specifically, tape-stripping was used to transiently disrupt the epidermal barrier, permitting the epicutaneous application of B16 melanoma cells at a single site per mouse. Within 2-6 weeks, ∼20-30% of mice displayed pigmented tumours at the site of inoculation, although some developed tumours as late as 6–14 weeks. Some such tumours were initially “controlled”, but essentially all progressed thereafter (Fig 6a). Nonetheless, many mice failed to develop tumours: a “non-developer” (ND) phenotype that remained free of macroscopic skin lesions long after epicutaneous inoculation (Fig. 6a). The ND phenotype was shown to reflect T_RM_-mediated regulation, consistent with which, a conspicuous T_RM_ compartment expressing elevated levels of CD103, CD69 and CD44 could be found at the inoculation sites in ND mice (Fig 6a, b), as well as in peritumoral areas in cases where tumours did develop (Fig 6c). Consistent with the importance of T_RM_ cells in regulating tumour development, both the number and size of tumours were increased in mice treated with FYT720 (Supp Fig 6a), that blocks the egress of primed αβ T cells from lymphoid organs to form T_RM_ compartments in the tissues. There was no significant difference between tumour incidence in FYT720-treated WT and TCRδ^−/-^ mice (Supp Fig 6a); hence, FYT720-induced tumour susceptibility cannot be attributed to any actions of DETC, consistent with previous reports (Scharschmidt et al., 2015).

**Figure 6.**
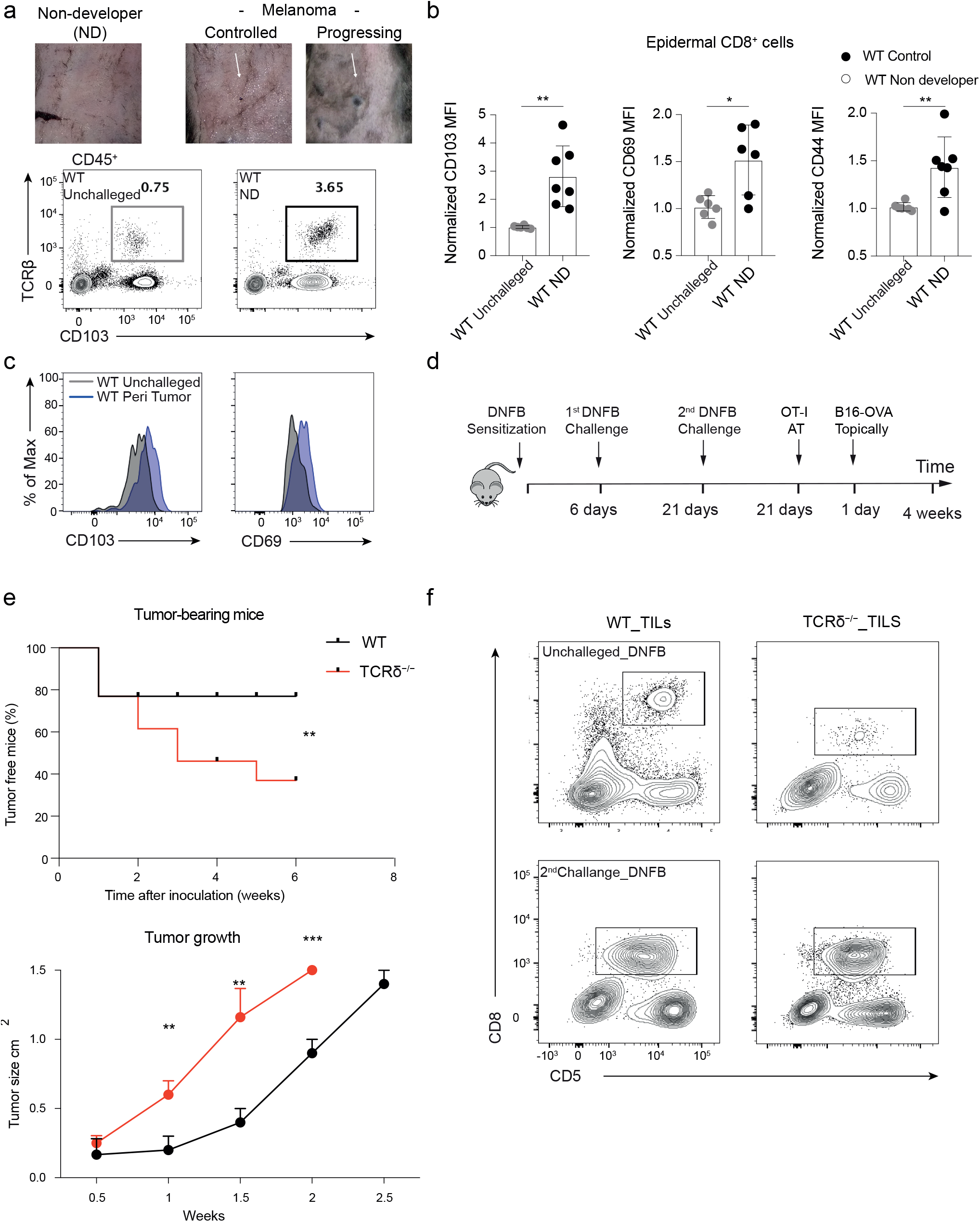
γδ T cell-deficiency impairs T_RM_-dependent tumour control. **a)** Macroscopic outcomes ≃30 days after epicutaneous inoculation of B16.OVA tumour cells (upper panels). Representative flow cytometry analysis of epidermal CD45^+^ cells in WT control *vs* WT ND mice (harvested from the site of inoculation) at ≃ weeks post inoculation (lower panels). **b)** Flow cytometric expression of indicated markers by epidermal CD8^+^ cells from WT control *vs* ND mice (harvested from the site of inoculation) at ≥3 weeks post inoculation. Data pooled from n = 2 biologically independent experiments with a total of 7 mice. Data are mean□±□SD. Statistical analysis was performed using Student’s t-test. ***p < 0.001, **p < 0.01, *p < 0.05. **c)** CD8^+^ phenotype in peritumoral skin at ≥3 weeks post inoculation, WT Control *vs* WT peri tumour mice. Representative CD103 and CD69 expression on CD8 cells isolated from the epidermis at ≥3 weeks B16.OVA inoculation. **d)** Workflow for adoptive transfer of CD8^+^ OT-I cells and B16.OVA inoculation after T_RM_ generation. **e)** Proportion of tumour-free WT *vs* TCRδ^−/−^ mice: statistical analysis performed using Kaplan-Meier test. ***p < 0.001, **p < 0.01, *p < 0.05. Data pooled from n = 3 biologically independent experiments with a total of 60 mice (upper panel). Tumour growth after B16.OVA inoculation in WT and TCRδ^−/−^ mice, n=10 (bottom panel). Data are mean□±□SD of a representative experiment of two independent experiments. Statistical analysis was performed using Student’s t-test. ***p < 0.001, **p < 0.01, *p < 0.05**. f)** Representative CD8 and CD5 expression assessed by flow cytometry on CD45^+^ tumour infiltrating cells from WT and TCRδ^−/−^ mice. ND: non-developer.

We then investigated melanoma formation in mice harbouring pre-formed T_RM_ compartments, a setting which probably better models the development of carcinomas in humans. Thus, we epicutaneously applied melanoma cells to mice that had been sensitised and twice-challenged over three weeks beforehand. Additionally, to recapitulate the development of some tumour-specific T cells, we used an ovalbumin (OVA)-expressing line of B16 melanoma, and supplemented the mice with an adoptive inoculum of OVA-reactive OT-1 cells (Fig 6d). As expected, the peritumoral areas in twice-challenged mice harboured substantive polyclonal T_RM_ compartments (compare Supp Fig 6c [top-row plots] with Fig 6c). Moreover, when the study was run in parallel in γδ T cell-deficient mice, the peritumoral areas harboured larger numbers of T_RM_ than did equivalent areas of WT mice, but the CD5 mean fluorescence intensity was slightly lower (Supp Fig 6c), completely consistent with the loss of DETC-mediated T_RM_ regulation. Indeed, we have frequently noted that T_RM_ generated in the absence of γδ T cells can also express lower levels of CD8, as shown here. Despite CD8^+^ Vα2^+^ OT1 cells being detected in the lymph nodes and spleen (Supp Fig 6d), they showed no appreciable peritumoral accumulation in either WT or TCRδ^−/−^ mice (Supp Fig 6c), possibly reflecting competition from substantial pre-accumulations of ACD-induced CD8^+^ T cells.

Rather than improving control over tumour formation, the enhanced frequency and effector phenotype of T_RM_ induced in TCRδ^−/−^ mice was associated with a decreased frequency of ND. Compared to ∼80% NDs on the WT background, there were <40% NDs on the TCRδ^−/−^ background, and those tumours that developed showed accelerated growth (Fig 6e). Strikingly, those tumours that developed in TCRδ^−/-^ mice were also frequently rich in CD8^+^ TILs (Fig 6f), but these were clearly insufficient to control tumour growth. Thus, in the setting of γδ deficiency, a tumour that is ordinarily limited by local T_RM_ became surrounded by a larger but qualitatively distinct T_RM_ compartment, and was less susceptible to host control. It is important to note that in this setting, the differential phenotype was not easily attributable to issues such as reduced barrier integrity in TCRδ^−/−^ mice, because based on trans-epidermal water loss, the impairment of barrier function which was induced by tape-stripping in this protocol was comparable in WT and TCRδ^−/−^ mice (Supp Fig 6d).

## Discussion

T_RM_ cells are a striking example of the critical role played by local immune systems in responding to secondary challenges wrought by allergens, reinfections, cancer recurrence, and even primary insults to which T_RM_ may be cross-reactive and/or make innate-like responses (Schenkel et al., 2014). It is therefore not surprising that they would be locally regulated. Thus, skin T_RM_ cells are known to be influenced by keratinocytes (Hirai et al., 2019) and there is likewise regulation of T_RM_ cells by T_reg_ cells that can be found in many extralymphoid tissues (Ferreira et al., 2020). Nonetheless, this study establishes that local regulation of T_RM_ cells is also profoundly mediated by local innate-like lymphocytes that become tissue-intrinsic during development, rather than following infection and/or other challenges, and that thereby create the environment into which T_RM_ cells will enter. Thus, in the absence of DETC, there was an exaggerated expansion of skin T_RM_ cells but they had an atypical phenotype including reduced CD5 expression that has been associated with lower intrinsic TCR affinity for antigen (Mandl et al., 2013). This result was observed in the setting of ACD, induced by a stimulus reflective of common human environmental exposures, and the results obtained justify *a posteriori* the value of investigating the impact of DETC on a polyclonal T_RM_ compartment, rather than on a transgenic compartment of uniform affinity. Analogous DETC regulation of T_RM_ maturation was also observed in mice challenged with PDV and with epicutaneous inocula of B16, respectively.

Innate-like lymphocytes including DETC are commonly viewed as protecting tissue integrity, promoting repair, and limiting inflammation (Fan and Rudensky, 2016; Hayday, 2019), but there has been surprisingly little consideration of whether their biology might include regulating the establishment and maturation of T_RM._ That these two qualitatively distinct tissue-associated lymphocyte universes are integrated has many implications. First, whereas DETC have been shown to make way for expanding T_RM_ at sites of challenge (Zaid et al., 2014), the near-equivalent de-repression of T_RM_ expansion in three different types of DETC-deficient mice cannot simply be attributed to their being more space for T_RM_ because DETC-deficient mice all harbour intraepidermal “replacement DETC”; moreover, normal numbers of DETC are present in mice with γδ-specific deletion of IFNγR2 in which T_RM_ expansion was also abnormally high. Indeed, it seems striking that T_RM_ maturation is dysregulated by the conditional ablation of a single gene in an heterologous cell type. Thus, DETC can be viewed as *bona fide* immunoregulators of T_RM_ cells, and the same likely holds for other tissue-intrinsic γδ T cells and innate-like lymphocytes. In this regard, we note that DETC express several molecules, including GITR, that have been described as properties of highly potent regulatory cells (Wyss et al., 2016), and likewise that TCRγδ^+^ intestinal IEL, which are conserved in humans, also express PD-L1 and several other molecules implicated in immunoregulation (Shires et al., 2001).

Second, we show that PD-L1 expression is at least one part of the mechanism by which DETC respond to IFNγ, and that antibody-mediated PD-L1 ablation phencopied major impacts of DETC on T_RM_ including the regluation of their motility which will be a key trait for cells operating within an anatomically confined space. Indeed, the data evoke two other highly disparate examples of PD-L1 regulation of T cell motility, observed in Type I diabetes (Fife et al., 2009) and in oncogene-induced tissue regeneration (Kortlever et al., 2017). In both cases, the aggregate outcome may be to limit T cell activation to only those T cells with sufficiently high TCR affinity to make strong-stops on local target cells. High affinity T cells are typically those reactive to foreign antigen by comparison to autoreactive T cells that escape cental tolerance because of reduced affinity (Cho and Sprent, 2018).

In this regard, we note that in mice adoptively transferred with T cells with lower intrinsic TCR affinity for influenza viruses, the lung T_RM_ compartment that developed in influenza-infected mice was very small compared to that which formed in mice receiving higher affinity cells, which also expressed higher levels of CD5 (Fiege et al., 2019). Thus, by integrating those data with our own, we can hypothesise that tissue-intrinsic γδ T cells impose filters that bias T_RM_ expansion toward motile, higher affinity CD5^hi^ cells, thereby minimising the prospect of T_RM_ compartments becoming over-populated with less effective and potentially autoreactive T cells. Consistent with this, removing these filters in multiple strains of γδ-deficient and IFNγR-deficient mice leads to auto-inflammatory, αβ T cell-dependent disease (Chen et al., 2002; Girardi et al., 2002; Mukasa et al., 1995).

Our findings evoke seminal studies of Amigorena and colleagues who showed that a key consequence of T_reg_ activity in *Listeria* infection was to limit memory compartments to higher affinity, more efficacious, anti-bacterial T cells that were less susceptible to T_reg_ control (Pace et al., 2012). In this light, it is also the case that the perpetuation of T_RM_ cells which are cross-reactive toward neo-antigens, for example, T_RM_ induced by common cold coronaviruses that have low affinity for SARS-CoV-2 antigens, may contribute to immunopathology (Bacher et al., 2020), and may offer another rationale for locally regulating T_RM_ quality.

Note however that this consideration does not dismiss the prospect that T cells bearing lower affinity TCRs may with relatively high efficiency form T_RM_ that, *via* cross-reactivity, provide valuable contributions to a first line of defence (Fiege et al., 2019; Lipsitch et al., 2020). Such contributions might be more important at later time points when the immediate prospects of re-encountering the initial challenge have declined. Possibly consistent with this perspective, T_RM_ cells with lower affinity displayed a gene expression profile that included *Bcl2* and other genes associated with cell survival (Fiege et al., 2019). Furthermore, once an epidermal T_RM_ compartment has become fully established, the regulatory influence of DETC will presumably be reduced by their evident spatial exclusion (Zaid et al., 2014). That is to say, local tissue-intrinsic regulation over T_RM_ quality is imposed early, seemingly consistent with our observations *in vitro* that DETC may regulate the maturation of T_RM_ progenitors.

Third, our studies strongly suggest that CD5 may be a useful discriminator of the quality of the T_RM_ compartment, as may be RANK-L. Although the peritumoral areas of TCRδ^−/−^ mice were well populated by large numbers of T_RM_, their aggregate lower expression of CD5 indicated that they will share the atypical cell surface, molecular, and cell biological phenotypes of polyclonal T_RM_ that we have shown to develop following ACD of γδ-deficient mice. Those T_RM_ were unable to assert control of local melanoma growth in the setting of γδ deficiency. Such observations illustrate that host-beneficial immunological responses to cancer, that are known to be multipartite (Plaks et al., 2015), may frequently involve the integration of one distinct lymphocytic lineage with another. Indeed, in another setting, adoptive transfer of CD8^+^ TIL that lacked Runx3 expression and which did not exhibit a signature T_RM_ phenotype failed to control tumour growth, by comparison to effective tumour control exerted by CD8^+^ T_RM_ overexpressing Runx3 (Milner et al., 2017). Additionally, T_RM_ infiltration has to date been shown by at least two independent studies to be a better prognostic marker of cancer suppression than was quantification of circulating CD8^+^ T cells (Ganesan et al., 2017; Nizard et al., 2017).

Fourth, we would propose that T_RM_ regulation by tissue-intrinsic innate-like lymphocytes that show many anatomic site-specific adaptations will provide a means to condition the responsiveness and effector potentials of T_RM_ to the physiology of particular tissues in defined species. Thus, site-specific variations in the responsiveness and effector potentials of T_RM_ cells may map back to variations in tissue-intrinsic innate-like lymphocytes at steady-state and in their responsiveness to local changes in tissue status. Indeed, T_RM_ regulation by tissue-intrinsic innate-like lymphocytes that sense the homeostatic *versus* stressed states of a tissue can provide T_RM_ with information that critically contextualises their exposure to antigen. Such considerations highlight the complexities as well as opportunities available to immunotherapeutics, with this study offering a case-in-point of an hitherto under-explored axis of T_RM_ regulation (motility) that may be directly influenced by the PD1/PD-L1 checkpoint. Such axes may improve our application of checkpoint blockade in sites such as the human bowel that harbour large compartments of tissue-intrinsic, innate-like T cells.

Finally, our findings seem germane to many instances of inflammatory pathologies associated with gd T cell deficiencies (Hayday and Tigelaar, 2003). Various aetiologies have been considered to underpin such disease states, including dysbiosis caused by a deficiency in dermal IL-17-producing gd T cells (Spidale et al., 2020). Notwithstanding such causes, tissue-associated ab T cells are a key effector of the described pathologies (Girardi et al., 2002; Shiohara et al., 1996), and this will clearly be exaggerated if local gd T cells are unable to regulate the quality and size of the T_RM_ compartment. One might therefore consider that enhancing local ecological regulation by tissue-intrinsic lymphocytes might ameliorate some examples of organ-specific autoimmune disease.

## Acknowledgements

We thank the Immunosurveillance laboratory members, particularly Leticia Monin, Flor Cano, Adam Laing, Shraddha Kamdar, and Natalie Roberts for advice and discussions; Anna Baulies (Francis Crick Institute) for data analysis; Sofia Mensurado for sharing protocols; and Bruno Silva-Santos and Francesca Di Rosa for discussions. Work was supported by: the Francis Crick Institute which receives its core funding from Cancer Research UK (CRUK), the UK Medical Research Council, and the Wellcome Trus (FC001003); the CRUK King’s Cancer Centre; the NIHR BRC at Guy’s and St Thomas’ NHS Foundation Trust and King’s College London; and the European Molecular Biology Organization (ALTF 198–2018 to MMR). MLI was supported in part by Gamma Delta Therapeutics. The funders had no role in study design, data collection and analysis, decision to publish, or preparation of the manuscript.

## Author contributions

ACH and MMR designed the study and experiments, MMR, AP, AMM, MLI, AJ, DU, BG-C and RD performed the experiments. MMR, DU, AJ, ML, DMc, and ACH processed and interpreted data. ACH wrote the manuscript; MMR, DMc, and ACH edited the text; MMR prepared figures.

---

**Supplemental Figure 1.**
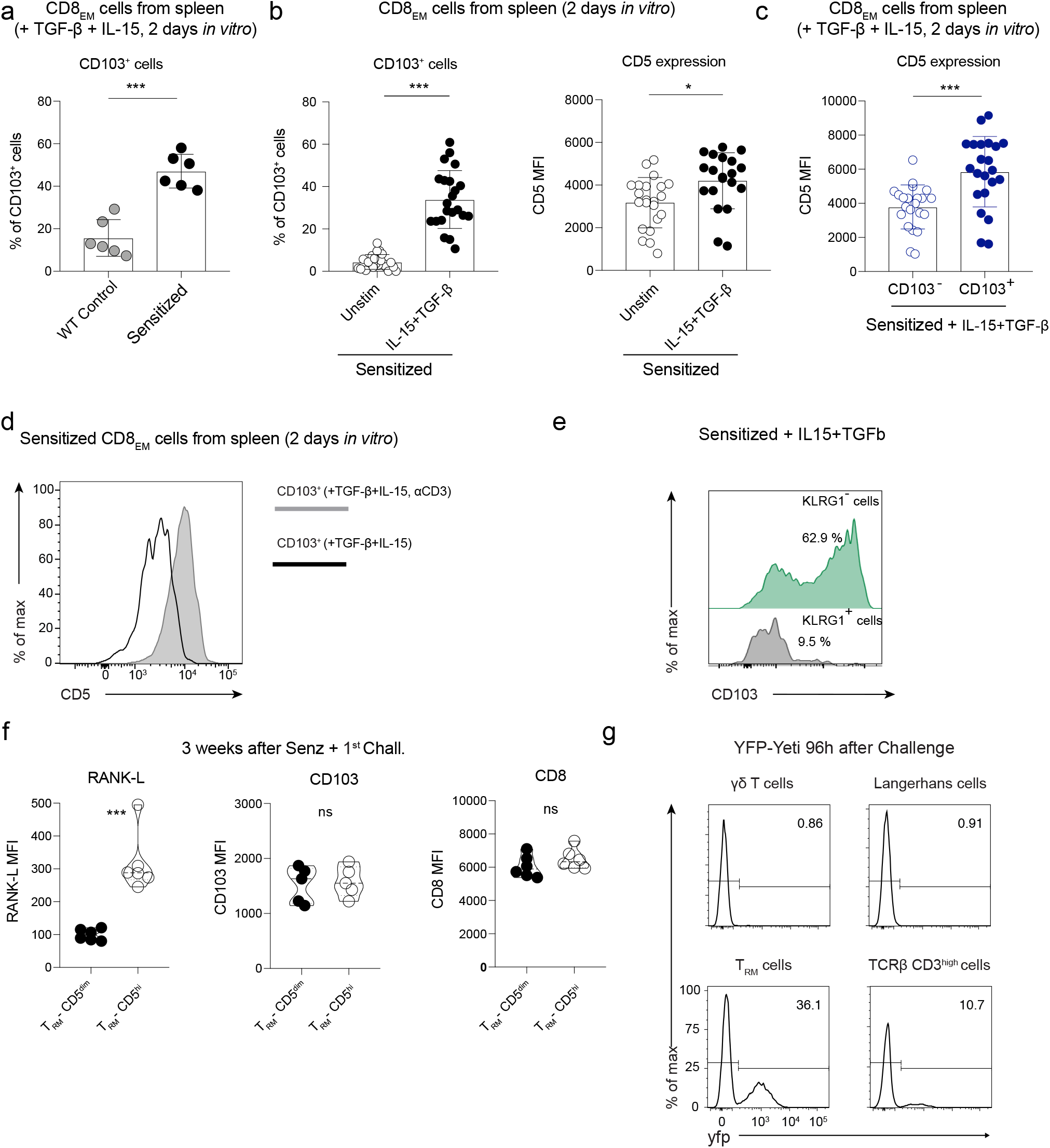
γδ T cell numbers critically regulate polyclonal T_RM_ generation and expansion, Related to Figure 1. **a-e)** Mice were senzatized with DNFB and spleen cells were harvest at d5 and cultured for 2d with TGF-b and IL-15. **a)** Frequency of conversion of CD8^+^ T cells, into T_RM_-like cells. **b)** Frequency of and CD5 expression by CD103^+^ CD8_EM_ cells. **c)** CD5 expression on CD103^+^CD8_EM_ *vs* CD103^−^CD8_EM_ cells (**a-c,** n = 20). **d)** CD5 expression on CD103^+^CD8_EM_ cells isolated from spleen 5 days after DNFB sensitization and cultured for 2 days with TGF-β and IL-15 or TGF-β, IL-15 and αCD3. Representative of 5 experiments. **e)** Representative plot of CD103 expression by KLRG1^+^ or KLRG1^−^ CD8_EM_ cells. **f)** Expression of RANK-L, CD103, or CD8 by CD8^+^CD5^dim^ and CD8^+^CD5^high^ cells 21 days after Sensitisation and 1^st^ Challenge in WT mice (n=5). Data are mean□±□SD. Statistical analysis was performed using Student’s t-test. ***p < 0.001, **p < 0.01, *p < 0.05**. g)** Representative analysis of yfp as a read-out of IFNγ was measured by flow cytometry 96h after 1^st^ Challenge for indicated populations of CD45^+^ epidermal cells in YFP-Yeti mice.

**Supplemental Figure 2.**
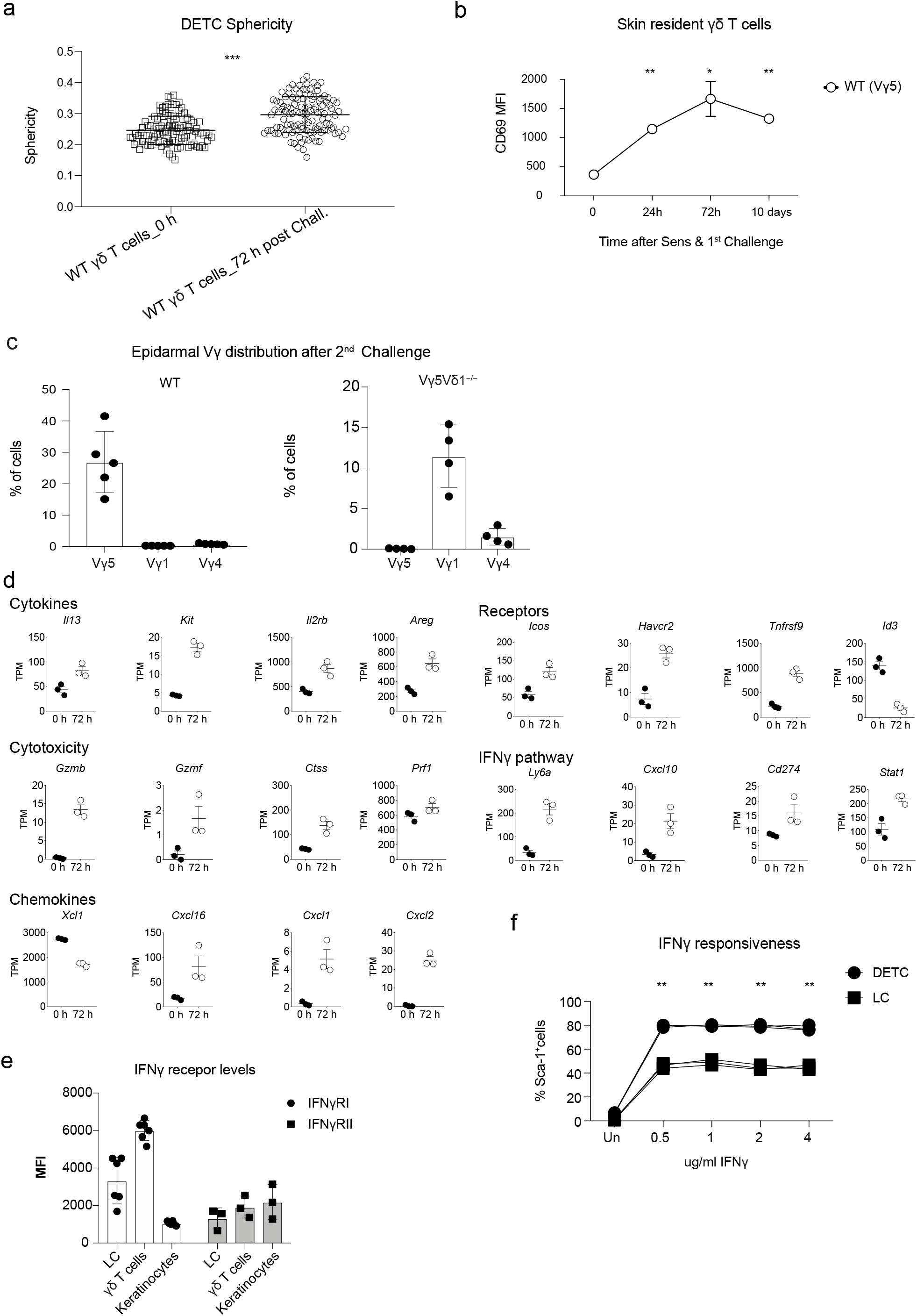
γδ T cell responsiveness to ACD and regulation by IFNγ. Related to Figure 2. **a)** Sphericity of DETCs by confocal microscopy at indicated times following DNFB challenge. **b)** CD69 was measured by flow cytometry at different time points after Sensitisation and 1^st^ Challenge on WT epidermal γδ T cells (n=5). **c)** Epidermal Vγ usage by gd T cells from WT and Vγ5Vδ1^−/−^ mcie 21 days after Sensitisation and 2^nd^ Challenge (nWT=5, nVγ5Vδ1^−/−^=4). **d)** RNAseq of sorted γδ T cells from unchallenged and challenged (72h after 1^st^ Challenge) WT mice. Individual data points are presented for different group of genes (n=3 n’=3.) **e)** IFNγRI and IFNγRII expression by flow cytometry in WT epidermis at steady-state (n=6). **f)** Comparison of IFNγ responsiveness by Sca-1 expression in WT DETC and LC after O/N stimulation with different concentrations of rIFNγ. n=5. Representative experiment of 2 independent experiments.

**Supplemental Figure 3.**
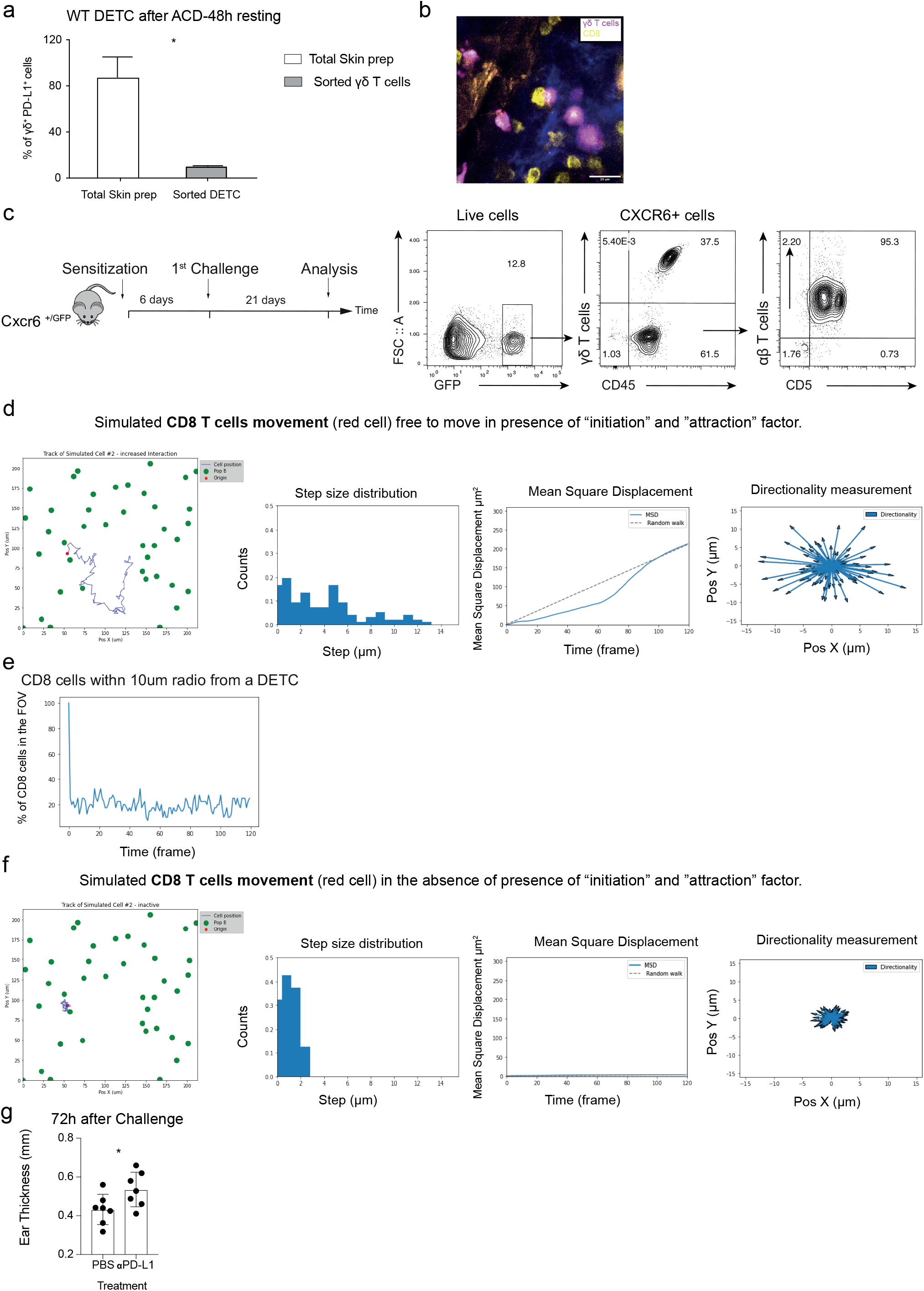
Local DETC-T_RM_ checkpoint implying direct contact, Related to Figure 3. **a)** Frequency of PD-L1^+^ DETC by flow cytometry on WT γδ T cells either alone or maintained in the presence of all other epidermal cells. Data are mean□±□SD of a representative experiment of 2 independent experiments. Statistical analysis was performed using Student’s t-test. *p < 0.05. **b)** Confocal microscopy of ear epidermal sheets of WT mice 72h after Sensitisation and 1^st^ Challenge. DETC stained for γδ TCR (purple) and CD8 T cells for CD8α (yellow). Scale bar as indicated in each image (µm). **c)** ACD experimental protocol in CXCR6^GFP/+^ mice and analysis of GFP expression on epidermal cells. Representative of 3 independent experiments. **d)** T_RM_ movement modelling. Representative images of: cell tracking or step size distribution or Mean Square Displacement or directionality “Simulated CD8 T cells movement (red cell) free to move in presence of “initiation” and “attraction” factor”. **e)** Modelling of 40 CD8 T cells movement in a given field-of-view containing sessile DETC. **f)** T_RM_ movement modelling. Representative images of: cell tracking or step size distribution or Mean Square Displacement or directionality “Simulated CD8 T cells movement (red cell) in the absence of presence of “initiation” and “attraction” factor”. **g)** Changes in ear thickness after Sensitization plus Challenge in WT mice treated with PBS or anti-PD-L1 (n = 7); ear thickness was measured at 72h after Challenge).

**Supplemental Figure 4.**
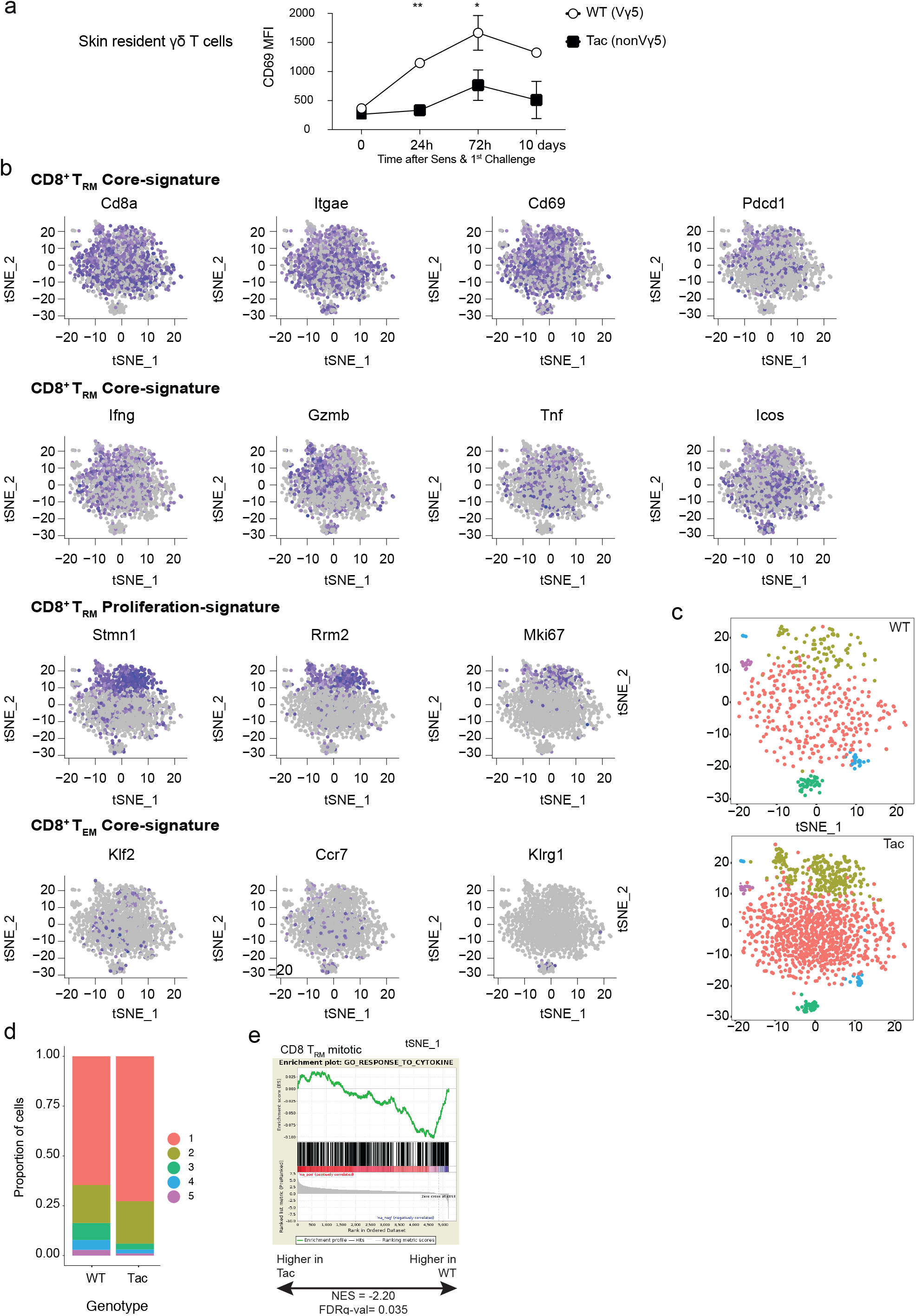
Altered molecular phenotypes of T_RM_ in γδ T cell-deficient mice, Related to Figure 4. **a)** CD69 expression in epidermal gd T cells by flow cytometry at indicated times after Sensitisation and 1^st^ Challenge of WT and Tac mice (n=5). **b)** Feature plots demonstrating expression of designated genes in the 1829 cells. **c)** t-SNE plot from Figure 3 segregated by genotype (WT *vs* Tac) in relation to 5 distinct clusters based on gene expression differences demarcated by colours. **d)** Percentages of cells from each cluster in as recovered from WT *vs* Tac mice. **e)** Gene set enrichment analysis (GSEA) for T_RM_ from Tac *vs* WT backgrounds using publicly available gene set. The enrichment score (NES) and p value are reported.

**Supplemental Figure 5.**
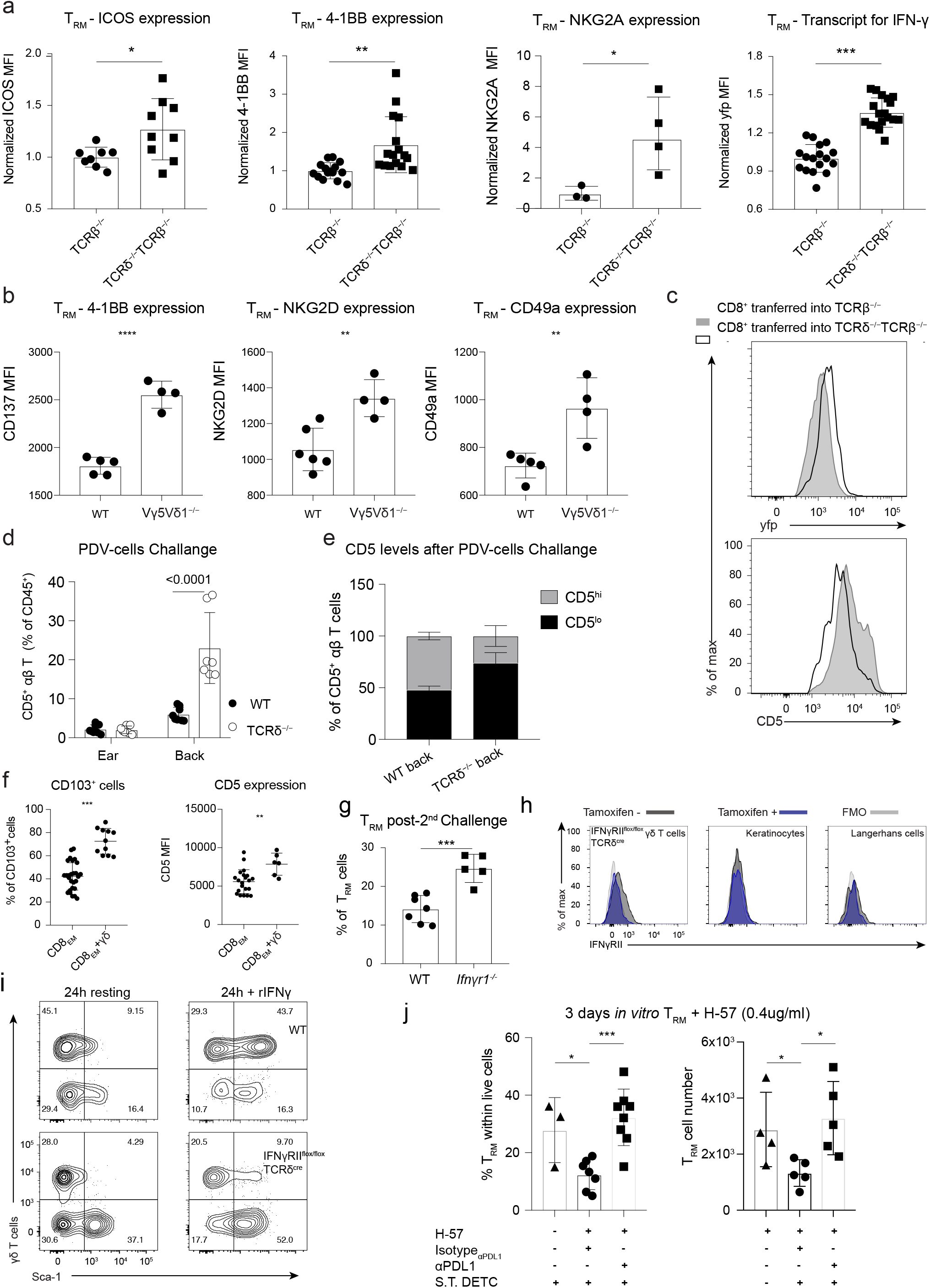
The absence of γδ T cells dysregulates T_RM_ homeostasis in the skin, Related to Figure 5. **a)** Comparative flow cytometric expression of indicated activation markers on epidermal CD8^+^ cells from YETI mouse following adoptive transfer to TCRβ^−/−^ or TCRδ^−/−^TCRβ^−/−^mice that were then sensitised, challenged, and analysed after three weeks. Normalized T_RM_ expression of yfp on CD8^+^. Data pooled from 3 biologically independent experiments with a total of 33 mice. Data are mean□±□SD. **b)** Expression of CD137, NKG2d, or CD49a of CD8^+^CD5^+^ cells 21 days after Sensitisation and 2^nd^ Challenge in WT and Vγ5Vδ^−/−^ mice (n=5). **c)** Flow cytometry analysis of CD5 expression by *i.v.* transferred CD8^+^ cells 21 days after Sensitisation and 1^st^ Challenge on TCRβ^−/−^ (grey histogram) and TCRδ^−/−^TCRβ^−/−^ (open histogram) backgrounds. **d)** T_RM_ numbers 12 weeks after inoculation of 10^6^ PDV cancer cell line (back) and resting skin (ear), expressed as CD5^+^ CD8 T cells in WT and TCRδ^−/−^ mice. **e)** CD5 levels after PDV-cells challenge in WT and TCRδ^−/−^ mice at the back (inoculation area). representative of two experiments **f)** Expression of CD103 and CD5 by sensitized CD8_EM_ cells after 3 days *in vitro* co-culture (CD8 + Short term DETC) with TGF-β and IL-15. **g)** Frequency of TRM amongst epidermal T cells 21 days after the 2^nd^ Challenge in WT and *Ifn*γ*r1^−/−^* animals, n=5. Data are mean□±□SD of two independent experiments. **h)** Flow cytometry analysis of IFNγRII expression in the indicated populations from *Tcrd^creER^Ifn*γ*r2^fl/fl^*mice following 4OH-tamoxifen treatment or vehicle (n=3). **i)** Comparison of IFNγ responsiveness (O/N stimulation with rIFNγ) by Sca-1 expression in WT DETC and *Tcrd^creER^Ifn*γ*r2^fl/fl^*mice following 4OH-tamoxifen treatment and ACD induction after (n=3). **j)** Frequency and number of T_RM_ after 3 days *in vitro* co-culture (2^nd^ Challenge T_RM_ + Short term DETC + H-57) performed in the presence of αPDL1 blocking antibody (20ug/ml) or isotype control (20ug/ml). Statistical analysis was performed using using one-way ANOVA with Sidak’s multiple comparisons post-hoc test**. a-h)** Statistical analysis was performed using Student’s t-test. ***p < 0.001, **p < 0.01, *p < 0.05.

**Supplemental Figure 6.**
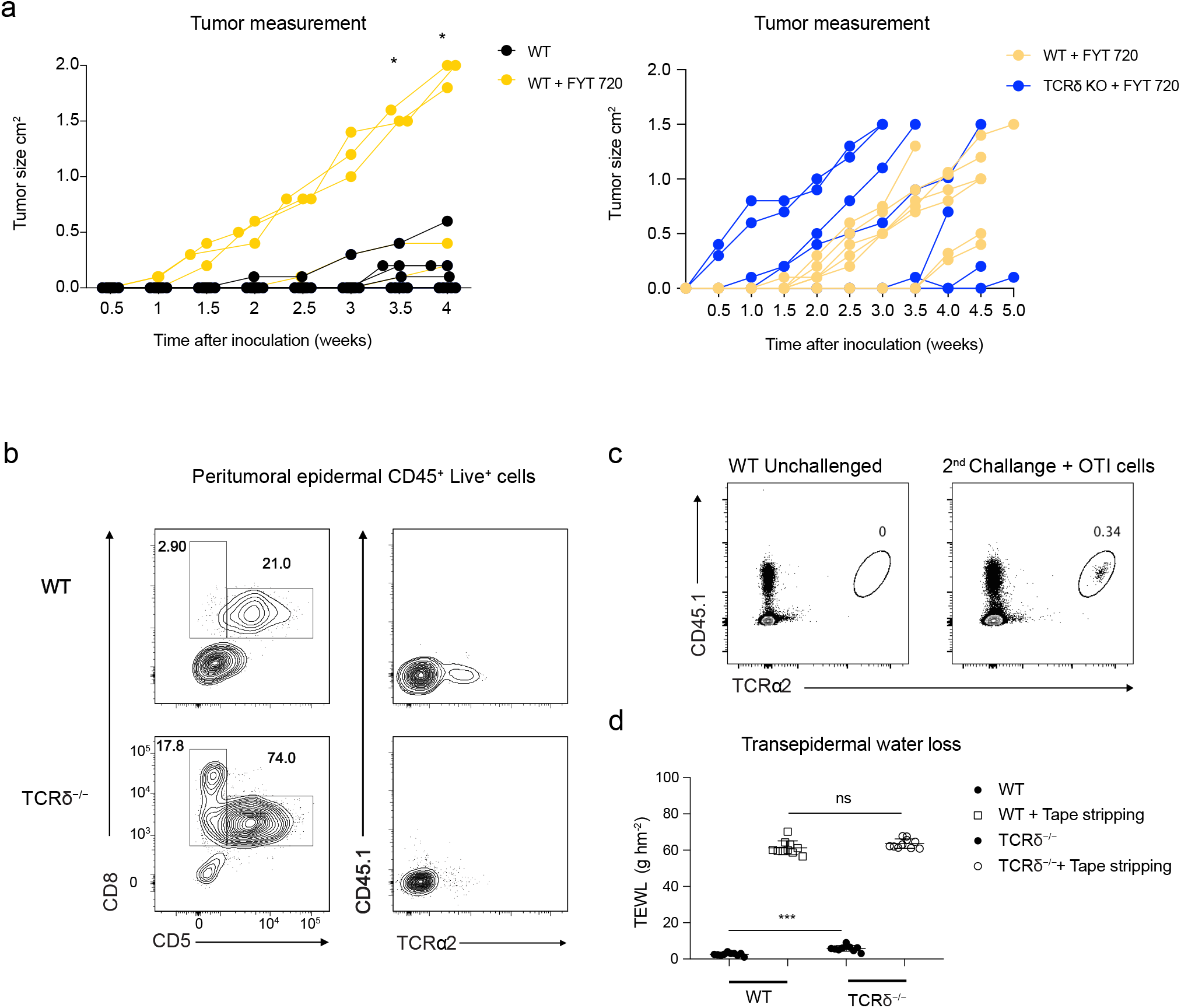
γδ T cells promote immune-mediated tumour control in a T_RM_-dependent model of transplantable epicutaneous melanoma, Related to Figure 6. **a)** Tumour growth after B16.OVA inoculation in WT mice with and without FYT720 treatment. Total of 20 mice (left panel). Tumour growth after B16.OVA inoculation in WT and *Tcrd*^−/−^ mice with FYT720 treatment. Total of 20 mice (right panel). Statistical analysis was performed using Student’s t-test of raw means with SD. ***p < 0.001, **p < 0.01, *p < 0.05. **b)** Representative plot of CD8 and CD5 expression assessed by flow cytometry on tumour infiltrating CD45^+^ cells (left panel). Staining for CD45.1 and TCRα2 was used to identify OT1 cells at 3 weeks post-adoptive transfer (Fig 6d) (right panel). **c)** Representative plot of CD45.1 and TCRα2 staining used to identify OT1 cells at 3 weeks post-adoptive transfer in the spleen. **d)** The dorsal skin was abraded by tape-stripping and trans-epidermal water loss (TEWL) measured prior to and immediately after tape-stripping using a tewameter probe assessing water evaporation rate and reported as g hm−2 (where g=water loss in grams, h=time in hours, m2=metres squared). TEWL was measured in WT and TCRδ^−/−^ mice. n=10 per group. Data are mean□±□SD. Statistical analysis was performed using Student’s t test. ***p < 0.001, **p < 0.01, *p < 0.05.

