## Supplementary material for "Tissue-intrinsic γδ T cells critically regulate Tissue-Resident Memory CD8 T cells": Materials & Methods

**STAR★METHODS**

**KEY RESOURCES TABLE**

| **REAGENT or RESOURCE** | **SOURCE** | **IDENTIFIER** |
| --- | --- | --- |
| Antibodies | | |
| Brilliant Violet 605 anti-mouse CD8a (53-6.7) | BioLegend | Cat# 100744, RRID: AB_2562609 |
| PE anti-mouse CD5 (53-7.3) | bdbiosciences | Cat# 553022, RRID:  AB_394560 |
| Brilliant Violet 605 anti-mouse CD5 (53-7.3) | bdbiosciences | Cat# 563194, RRID:  AB_2738061 |
| FITC anti-mouse Ly-6A/E (Sca-1) (D7) | eBioscience | Cat# 11-5981-82, RRID:  AB_465333 |
| APC anti-mouse CD103 (2E7) | BioLegend | Cat# 121413, RRID:  AB_1227503 |
| PE anti-mouse CD103 (2E8) | BioLegend | Cat# 121406, RRID:  AB_1133989 |
| PE/Cy7 anti-mouse CD8a (53-6.7) | BioLegend | Cat# 100722, RRID:  AB_312761 |
| Pacific Blue anti-mouse CD8a (53-6.7) | BioLegend | Cat# 100725, RRID:  AB_493425 |
| PE/Cy7 anti-mouse CD69 (H1.2F3) | BioLegend | Cat# 104512, RRID:  AB_493564 |
| APC/Cyanine7 anti-mouse CD69 (H1.2F3) | BioLegend | Cat# 104526, RRID:  AB_10679041 |
| PE/Cy7 anti-mouse/human CD44 (IM7) | BioLegend | Cat# 103030, RRID:  AB_830787 |
| PerCP/Cyanine5.5 anti-mouse (30-F11) | BioLegend | Cat# 103132, RRID:  AB_893340 |
| APC/Cyanine7 anti-mouse CD45 (30-F11) | BioLegend | Cat# 103116, RRID:  AB_312981 |
| FITC anti-mouse TCR gamma/delta (GL3) | BioLegend | Cat# 118106, RRID:  AB_313830 |
| Brilliant Violet 421 anti-mouse TCR gamma/delta (GL3) | BioLegend | Cat# 118119, RRID:  AB_10896753 |
| Brilliant Violet 421anti-mouse TCR beta (H-57) | BioLegend | Cat# 109230, RRID:  AB_2562562 |
| APC anti-mouse TCR beta (H-57) | BioLegend | Cat# 109212, RRID:  AB_313435 |
| Brilliant Violet 605 anti-human/mouse/rat CD278 (ICOS) (C398.4A) | BioLegend | Cat# 313538, RRID:  AB_2687079 |
| Alexa Fluor 647 anti-human/mouse/rat CD278 (ICOS) (C398.4A) | BioLegend | Cat# 313516, RRID:  AB_2122582 |
| APC anti-mouse CD314 (NKG2D) (CX5) | BioLegend | Cat# 130212, RRID:  AB_1236372 |
| PE anti-mouse CD314 (NKG2D) (CX5) | BioLegend | Cat# 130207, RRID:  AB_1227713 |
| FITC anti-mouse CD314 (NKG2D) (C7) | BioLegend | Cat# 115711, RRID:  AB_2133291 |
| PE/Cy7 anti-mouse CD159a (NKG2AB6) (16A11) | BioLegend | Cat# 142809, RRID:  AB_2728160 |
| APC/Cyanine7 anti-mouse CD3epsilon (2C11) | BioLegend | Cat# 100330, RRID:  AB_1877170 |
| Brilliant Violet 605 anti-mouse CD119 (IFN-γ Rα) (GR20) | bdbiosciences | Cat# 745111, RRID:  AB_2742716 |
| PE IFN-γR β chain Antibody | Miltenyi Biotec | Cat# 130-105-671, RRID: AB_2652252 |
| Brilliant Violet 605 anti-mouse CD366 (Tim-3) (RMT3-23) | BioLegend | Cat# 119721, RRID:  AB_2616907 |
| PE anti-mouse CD254 (TRANCE, RANKL) (IK22/5) | BioLegend | Cat# 510005, RRID:  AB_315553 |
| PE anti-mouse CD137 antibody (17B5) | BioLegend | Cat# 106106, RRID:  AB_2287565 |
| FITC anti-mouse CD279 (PD-1) (29F.1A12) | BioLegend | Cat# 135214, RRID: AB_10680238 |
| Brilliant Violet 421 anti-mouse CD274 (B7-H1, PD-L1) (10F.9G2) | BioLegend | Cat# 124315, RRID: AB_10897097 |
| PE anti-mouse TIGIT Monoclonal Antibody (GIGD7) | eBioscience | Cat# 12-9501-82, RRID: AB_11042152 |
| Purified anti-mouse CD274 (B7-H1, PD-L1) (10F.9G2) | BioLegend | Cat# 124302, RRID: AB_961228 |
| Purified Rat IgG2b, κ Isotype control (RTK4530) | BioLegend | Cat# 400601, RRID: AB_326545 |
| TCR beta Monoclonal Antibody (H57-597), Functional Grade. | eBioscience | Cat# 16-5961-82, RRID: AB_469169 |
| Chemicals, Peptides, and Recombinant Proteins | | |
| Fetal Bovine Serum, Heat Inactivated | Gibco | Cat# 10270106 |
| Penicillin-Streptomycin | Sigma-Aldrich | Cat# P4333 |
| RPMI1640 | Gibco | Cat# 21875-091 |
| LIVE/DEAD Fixable Blue Dead Cell Stain Kit | Thermo Fisher ScientifiC | Cat# L34962 |
| Hoechst 33342 | Thermofisher Scientific | Cat# H3570 |
| Olive oil | Sigma-Aldrich | Cat# 75343 |
| Acetone | Thermo Scientific | Cat# |
| 1-Fluoro-2,4-dinitrobenzene (DNFB) | Sigma-Aldrich | Cat# D1529 |
| Red Blood Cell Lysing Buffer Hybri-Max | Sigma Aldrich | Cat# R7757 |
| 4-OH Tamoxifen | Sigma Aldrich | Cat# H7904 |
| Ammonium thiocyanate | Sigma Aldrich | Cat# A7149 |
| Trypsin from bovine pancreas | Sigma Aldrich | Cat# T1005 |
| DNase I from bovine pancreas | Sigma Aldrich | Cat# 11284932001 |
| Trypsin/EDTA solution | Gibco | Cat# 25200-056 |
| HBSS (Hank's Balanced Salt Solution) | Thermofisher Scientific | Cat# 14175095 |
| Collagenase type IV | Worthington | LS004188 |
| DNAse I | Sigma-Aldrich | Cat# D5025-15 |
| BSA | Sigma Aldrich | Cat# A4503 |
| rTGFb | PeproTech | Cat# 100-21 |
| rmIL-15 | ImmunoTools | Cat# 12340155 |
| Matrigel | Corning | Cat# 734-0270 |
| Critical Commercial Assays | | |
| CD8a+ T Cell Isolation Kit, mouse | Miltenyi Biotec | Cat# 130-104-075 |
| Deposited Data | | |
| RNA sequencing data | This paper | GEO: GSE63473 |
| scRNA sequencing data | This paper |  |
| Experimental Models: Cell Lines | | |
| B16.OVA | The Francis Crick Institute | N/A |
| Experimental Models: Organisms/Strains | | |
| Mouse: WT: C57BL/6J | The Jackson Laboratory | Cat# JAX:000664; RRID:  IMSR_JAX:000664 |
| Mouse: C57BL/6-Tg(Nr4a1-EGFP/cre)820Khog/J | The Jackson Laboratory | Cat# JAX:016617, RRID:  IMSR_JAX:016617 |
| Mouse: B6.129P2-Tcrdtm1Mom/J | The Jackson Laboratory | Cat# JAX:002120, RRID:IMSR_JAX:002120 |
| Mouse: B6.129P2-Tcrbtm1Mom/J | The Jackson Laboratory | Cat# JAX:002118, RRID:IMSR_JAX:002118 |
| Mouse: B6.129P2-Tcrbtm1Mom Tcrdtm1Mom/J | The Jackson Laboratory | Cat# JAX:002122, RRID:IMSR_JAX:002122 |
| Mouse: IFNγ-IRES-YFP-BGHpolyA knockin (YETI) | The Francis Crick Institute | Stetson et al., 2003 |
| Mouse: B6.129S7-Ifngr1tm1Agt/J | The Jackson Laboratory | Cat# JAX:003288, RRID:IMSR_JAX:003288 |
| Mouse: IFN-γR2 floxed | The Francis Crick Institute | (Lee et al., 2015) |
| Mouse: B6.129S-Tcrdtm1.1(cre/ERT2)Zhu/J | The Jackson Laboratory | Cat# JAX:031679, RRID:IMSR_JAX:031679 |
| Mouse: FVB/NJ (FVB.WT) | The Francis Crick Institute | Cat# JAX:001800, RRID:IMSR_JAX:001800 |
| Mouse: FVB.δ^−/−^ | The Francis Crick Institute |  |
| Mouse: FVB/NTac (FVB.Tac) | The Francis Crick Institute |  |
| Mouse: FVB.Vg5^−/−^;Vd1^−/−^ | The Francis Crick Institute |  |
| Mouse: FVB.β^−/−^ | The Francis Crick Institute |  |
| Mouse: FVB.β^−/−^δ^−/−^ | The Francis Crick Institute |  |
| Mouse: FVB.Cxcr6tm1Litt | The Francis Crick Institute |  |
| Mouse: FVB/N.R26 mTmg | The Francis Crick Institute |  |
| Software and Algorithms | | |
| ImageJ | Schneider et al., 2012 | <https://imagej.nih.gov/ij/>  RRID: SCR_003070 |
| GraphPad Prism versione 8.0.2 | Graphpad Inc | RRID: SCR_002798 |
| FlowJo version 10 | Treestar Inc | RRID: SCR_008520 |

**RESOURCE AVAILABILITY**

**Lead Contact**

**Materials Availability**

Mouse lines generated in this study will be maintained in the lead author’s current institute’s animal house and/or stored locally as frozen embryos and can be made available upon request.

**Data and Code Availability**

The RNA-seq datasets reported in this paper can be found at GEO: GSE164023 (GSE164022)

The scRNA-seq datasets reported in this paper can be found at GEO: GSE164023 (GSE164021)

The RNA-seq dataset reported in Fig 4.d has been submitted: GEO: GSE160477

**EXPERIMENTAL MODEL AND SUBJECT DETAILS**

**Mice**

Adult male mice were used at 4–20 weeks of age. All mice were bred at the Francis Crick Institute. C57BL/6J background: C57BL/6J.WT, TCRδ^−/−^, TCRβ^−/−^, TCRδ^−/−^; TCRβ^−/−^, YFP-enhanced transcript for IFN-γ (Yeti)(Stetson et al., 2003), Nur77^GFPCre^(Moran et al., 2011), *Ifnγr1*^–/–^ (Huang et al., 1993) and *Ifnγr2^fl/fl^* (Lee et al., 2015);Tcrd^Cre^(Zhang et al., 2015). *Ifnγr2^fl/fl^* mice were kindly provided by Jean Langhorne (The Francis Crick Institute, London UK). FVB background: FVB.WT, FVB.δ^−/−^, FVB.Vγ5^−/−^;Vδ1^−/−^, FVB.β^−/−^, FVB.β^−/−^δ^−/−^, FVB.*cxcr6*^tm1Litt^ and FVB/NTac (Tac) mice carrying a mutation in the *Skint1* gene were from Taconic farms. Mice were kept in filter-topped cages with sterilized food and water *ad libitum*, and autoclaved corncob bedding, changed at least once weekly. All experiments were performed according to the UK animal protection laws.

**METHOD DETAILS**

**Contact Dermatitis**

To induce allergic contact dermatitis (ACD), mice were sensitized on day 0 by epicutaneous application to razor-shaved abdominal skin (with a Wella razor and depilated using wecprep blades from Pilling) of 40 μL of 0.5% DNFB in a mixture of acetone:olive oil (3:1). On day 6, mice were challenged (1^st^ Challenge) by applying 20 μL of 0.25% DNFB in acetone:olive oil at the back skin or ear. In some cases, mice were rechallenged 21 days later (2^nd^ Challenge). In collaboration with Making STP at the Francis Crick Institute, a device was developed to assure the area of application was consistent across the experiments (50 mm^2^).

**Adoptive transfers**

For adoptive transfer, CD8^+^ T cells were purified from spleen and lymph nodes by CD8α^+^ T Cell Isolation Kit, mouse (Miltenyi Biotec) according to the manufacturer’s instructions and 1x10^6^ WT CD8^+^ T cells were intravenously transferred into T cell deficient animals denoted in the text. The following day, mice were sensitized and challenged as described above.

**Topical tamoxifen treatment**

4-Hydroxytamoxifen (4-OH tamoxifen) (H7904-25MG; Sigma-Aldrich) was dissolved in 99.5% acetone following incubation at 37°C for 5-10 min with occasional vortexing. A final concentration of 12 mg/ml was administered to mice for 5 consecutive days by topical application (20μl/ear).

**B16 melanoma inoculation**

For B16 melanoma inoculation, animals were placed individually in an induction chamber, and anesthetized with 5% isoflurane (Isoflo, Zoetis, UK) in 100% oxygen with a delivery rate of 5 L/min until loss of righting reflex. Lubricating eye gel (Lubrithal Eye Gel 10G Tube) was applied to the eyes to prevent drying. Mice were shaved (with a Wella razor) and depilated using wecprep blades (Pilling). B16 melanoma cells were harvested by washing with PBS, incubating cells at 37°C for 5 min with 1 × trypsin/EDTA solution (0.25% Trypsin-EDTA-Sigma), and washing with Hanks’ balanced saline solution (HBSS). For epicutaneous inoculation, the stratum corneum of back skin was removed by application and removal of cellophane tape (ScotchTM) five to eight times, and the scarified site was wiped with a cotton-tipped applicator soaked in PBS. B16.OVA cells (2×10^5^) were suspended in 20 μL of Matrigel basement membrane matrix (Corning) and applied to the scarified region. Mice were rested for about 10 min to allow solidification of Matrigel before application of Tegaderm film (3M) over the gel. It was removed 4 days later. Developing tumours were measured using a digital caliper.

**CD8 *in vitro* culture**

Total CD8^+^ T cells were isolated (CD8α^+^ T Cell Isolation Kit, mouse - Miltenyi Biotec) from spleens of sensitized mice 5 days after DNFB application and transferred into 96 well-plates with cytokines and/or antibodies described in the text. Two days after, CD8^+^ T cells were analyzed phenotypically by flow cytometry.

**Isolation of primary DETC lines**

Primary DETC were isolated and grown as previously described (Witherden et al., 2010). After approximately 1 month, cells were assessed for purity by flow cytometry. Cell lines with ≥85% DETC were used for coculture experiments.

**T_RM_-DETC coculture assays**

Primary DETC lines were isolated as described previously. T_RM_ were sorted (see below) 21 days after the 2^nd^ Challenge and added to primary DETC cultures in a 96-well plate at a concentration of 4 × 10^4^ cells/type of cell. All cells were collected 72h later.

**Flow Cytometry**

To analyze epidermal T cell populations, separate epidermal cell suspensions were prepared as described (Mallick-Wood et al., 1998) from ears or back skin of individual animals. When back skin was analyzed, always the same amount of skin (equivalent to 962mm^2^) was digested. After overnight culture to allow re-expression of trypsin-sensitive epitopes, epidermal cells were blocked with normal hamster IgG plus anti-FcR (2.4G2), and stained with anti-CD8α (53-6.7), anti-CD5 (53-7.3), anti-Ly-6A/E (Sca-1) (D7), anti-CD103 (2E7), anti-CD69 (H1.2F3), anti-CD44 (IM7), anti-CD45 (30-F11), anti-TCR gamma/delta (GL3), anti-TCR beta (H-57), anti-CD278 (ICOS) (C398.4A), anti-CD314 (NKG2D) (CX5), anti-CD314 (NKG2D) (C7), anti-CD159a (16A11), anti-CD3epsilon (2C11), anti-CD119 (GR20), anti-CD366 (RMT3-23), anti-CD254 (IK22/5), anti-CD137 antibody (17B5), anti-CD279 (PD-1) (29F.1A12), anti-CD274 (B7-H1, PD-L1) (10F.9G2) and anti-TIGIT (GIGD7). Isotype-matched control antibodies were used at the same concentrations as test antibodies. Analysis was performed with a FACScan™ (Becton Dickinson) with electronic gates set on live cells by a combination of forward and side light scatter and propidium iodide exclusion. A minimum of 100 live events was collected per sample and data were analyzed with FlowJo™ Software.

To analyze TILs, tumours were harvested, finely chopped, and digested with 1 mg/ml collagenase Type I, 0.4 mg/ml collagenase Type IV (Worthington), and 10 μg/ml DNase I (Sigma/Merck) for 30 minutes at 37 C with agitation, in medium without FBS. After this incubation, complete medium (with FBS) was added to stop the digestion reaction. The cell suspension and remaining tumors pieces were macerated in a 100μm cell strainer with a syringe plunger.

**FACS sorting for bulk RNA-seq and scRNA-seq**

To isolate epidermal T cell populations, separated epidermal cell suspensions were prepared as described from back skin of individual animals (Mallick-Wood et al., 1998). Incubation with monoclonal antibodies for 30 min at 4°C in FACS buffer (2% FBS in Phosphate Buffered Saline, pH 7.4, stabilized with 0.09% sodium azide), then washed twice in FACS buffer. Viable CD45^+^CD103^+^TCRβ^+^CD5^+^ for scRNA-seq and CD45^+^CD3^+^TCRδ^+^ for bulkRNA-seq cells were sorted into PBS buffer + 0.04% BSA and retained on ice. Sorted cells were confirmed to be >85–95% pure prior to RNA extraction.

**RNA sequencing**

cDNA was prepared from 10ng input RNA using the NuGEN Ovation RNA-Seq System (V2), and libraries constructed using the NuGEN Ultralow Library System V2.  Both these steps followed the manufacturer’s instructions.  The resulting libraries were pooled for sequencing on an Illumina HiSeq 2500 platform with single-ended 75 bp reads. Read adaptor removal and quality trimming was carried out with Trimmomatic (version 0.36)(Bolger et al., 2014). Reads were then aligned to the mouse genome, using Ensembl GRCm38 - release 86 as reference. Read alignment and gene level quantification was performed by STAR alignment (v.2.5.2a)(Dobin et al., 2013) together with RSEM package (v.1.2.31)(Li and Dewey, 2011). Differential expression analysis was carried out with DESEq2 (v 1.28.0)(Love et al., 2014) within R programming environment (R Core Team, 2017). Genes are called differentially expressed if padj < 0.05.

**Single-cell RNA sequencing.**

Samples were prepared using the 10x 3’ mRNA-Seq kit version 3.0. Briefly, cell viability was assessed using an EVE cell counter and Trypan blue viability stain. Approximately 10,000 cells were loaded in to the 10x Chromium which was operated according to manufacturer’s instructions. Sequencing was carried out on the Illumina HiSeq 4000 with a read configuration recommended by 10x for the sequencing of these libraries (28bp read 1, 98bp read 2, 8bp index 1). The 10X Cell Ranger software (version 2.1.1) was used to de-multiplex Illumina BCL output, create fastq files and generate single cell feature counts for each library using mm10-v1.2.0 as reference. All subsequent analyses were performed in R v.3.5.1 using the Seurat (v 2.3.4) package(Butler et al., 2018). Genes were removed if they were expressed in 3 or less cells and cells with less than 500 genes detected were also removed. Data was integrated following Seurat's vignette. In brief, for each sample the top 1000 most variable genes were selected for data integration using Canonical correlation Analysis (CCA) with 25 dimensions used for dimensional reduction using tSNE and cluster calling. Upon initial examination of the clusters, we excluded cells that expressed Cd47 more than 4 times (myeloid cells) and Krt15 more than 2 times (keratinocytes), prior integration and the analysis was repeated with the same parameters using resolution of 0.4 to define clusters. Cluster markers were identified using "FindConservedMarkers" with default parameters. To find genes expressed differentially between WT and Tac mice within each cluster, we used the DESeq2 test in "FindMarkers". Gene set enrichment analysis (GSEA) was carried out with the GSEA software (version 2.2.3) from the Broad Institute. The software compares ranked lists of genes, in this case, differentially expressed genes ranked by the "stat" value obtained from DESeq2 in decreasing order, with the following genesets "c2.cp.v7.0.symbols.gmt", "c5.bp.v7.0.symbols.gmt" downloaded from the Broad Institute.

**qRT-PCR**

Epidermis from back skin was separated from dermis as described in the flow cytometry section. To isolate specific epidermal T cell populations cell suspensions were prepared as described above in FACS sorting section**.** Samples were directly frozen in RLT buffer prior to RNA purification with DNAse digest (QIAGEN RNeasy kit). cDNA was generated using Superscript-II (Invitrogen) and analysed using Sybr-green assay (Invitrogen) using a Viaa7 Real-time PCR machine (Applied Biosystems). cDNA was analyzed for *Skint1* and *Skint2*, and expression levels of each gene normalized to Cyclophilin.

**Confocal microscopy**

Ears epidermis were separated with 0.5M Ammonium thiocyanate in PBS. The mechanically split ears floated with their dermal side down in the Petri dish and incubated in CO_2_ incubator at 37^o^C for 35 min. The epidermis was physically separated from the dermis and it was fixed with cold acetone for 20 min. The epidermal sheets were blocked with PBS containing 5% normal goat serum and stained for 1h with antibodies at RT in PBS containing 5% normal goat serum and 0.5% BSA usingTCRδ-BV421 and CD8-FITC antibodies. Z-Sections were acquired on a Leica SP5 confocal microscope using a 10x / 1.25 NA objective and processed and analyzed using either Fiji(Schindelin et al., 2012). For quantitative analysis, confocal records were processed to identify cells shown in Fig 4b using Otsu's automated thresholding and Multiresolution segmentation in Definiens Developer XD software. Morphological parameter sphericity was calculated as the ratio of cell object border length to its volume, ranging from 0 to 1 with higher values corresponding to more spherical objects.

**Ex-vivo confocal microscopy.**

Challenged ears of the animals were removed and placed on a #1.5 coverslip to be imaged on a Zeiss Upright LSM880 NLO, with a Plan-Apochromat 20x/0.8 NA objective. Cell’s migration was monitored exciting sequentially the sample with 488 nm and 561 nm laser lines, detecting the signal between 510-550 nm (GFP channel) and 580-640 nm (tdTomato channel). A volume of 213mm x 213mm x 30mm (z-step=5 mm) was acquired every 30 seconds (time interval), for a total duration of 30-60 minutes. Cells were manually tracked using the Manual Tracking plugin available in FIJI (Schindelin, J., 2012) and the tracks analysed (steps distribution, mean square displacement, and directionality) with custom made python scripts.

**T_RM_ movement modelling**

The simulation includes two cell populations, one set to be steady, with no change in XY position (green cells, imitating DETCs), the other allowed to move (red cells, imitating T_RM_ cells). The behavior of the simulated red cells has been modelled including two terms that change the possible movement, compared to randomly generated step size and angles: an “initiation” factor that increases the possible maximum step size, and an “attraction” factor that allows the cell to move with bigger steps towards the nearest cell of the second population, if the latter (green cell) happens to be within a proper distance (attraction radius).

Additionally, the model includes terms to simulate potential contacts between the two cell populations; in the case of close proximity with a green cell, the red cell direction is generated according to an inhomogeneous angle distribution.

The parameters used to simulate “1” were step_max=14 um, activation=0.99, attraction_radius=100 um, attractionProbability=0.7, while “2” has been simulated using step_max=14 um, activation=0.2, attraction_radius=100 um, attractionProbability=0.01.

**Statistical Analysis.**

Groups were compared with Prism software (GraphPad) using the two-tailed unpaired Student’s t test for comparison of two groups or the Kaplan Meier test to assess the probability of an event at a respective time interval (survival). Data are presented as each data point and mean or mean ± standard error of the mean (SD). p < 0.05 was considered significant. The experiments were not randomized and the investigators were not blinded to allocation during experiments and outcome assessment. No statistical methods were used to predetermine sample size. All experiments were performed at least twice, either with similar results obtained and representative data shown, or with pooled data shown. Survival curves and dotted graph bars with pooled data from multiple experiments include all experiments performed. Where representative histograms are shown, data reflect at least two independent experiments with at least three mice per experiment, in which similar results were obtained.

**BIBLIOGRAPHY (METHODS)**

Bolger, A.M., Lohse, M., and Usadel, B. (2014). Trimmomatic: a flexible trimmer for Illumina sequence data. Bioinformatics *30*, 2114–2120.

Butler, A., Hoffman, P., Smibert, P., Papalexi, E., and Satija, R. (2018). Integrating single-cell transcriptomic data across different conditions, technologies, and species. Nat. Biotechnol. *36*, 411–420.

Dobin, A., Davis, C.A., Schlesinger, F., Drenkow, J., Zaleski, C., Jha, S., Batut, P., Chaisson, M., and Gingeras, T.R. (2013). STAR: ultrafast universal RNA-seq aligner. Bioinformatics *29*, 15–21.

Huang, S., Hendriks, W., Althage, A., Hemmi, S., Bluethmann, H., Kamijo, R., Vilcek, J., Zinkernagel, R.M., and Aguet, M. (1993). Immune response in mice that lack the interferon-gamma receptor. Science *259*, 1742–1745.

Lee, H.-M., Fleige, A., Forman, R., Cho, S., Khan, A.A., Lin, L.-L., Nguyen, D.T., O’Hara-Hall, A., Yin, Z., Hunter, C.A., et al. (2015). IFNγ Signaling Endows DCs with the Capacity to Control Type I Inflammation during Parasitic Infection through Promoting T-bet+ Regulatory T Cells. PLoS Pathog. *11*, e1004635.

Li, B., and Dewey, C.N. (2011). RSEM: accurate transcript quantification from RNA-Seq data with or without a reference genome. BMC Bioinformatics *12*, 323.

Love, M.I., Huber, W., and Anders, S. (2014). Moderated estimation of fold change and dispersion for RNA-seq data with DESeq2. Genome Biol. *15*, 550.

R Core Team (2017). R: A language and environment for statistical computing. R Foundation for Statistical Computing, Vienna, Austria. URL https://www.R-project.org/.

Mallick-Wood, C.A., Lewis, J.M., Richie, L.I., Owen, M.J., Tigelaar, R.E., and Hayday, A.C. (1998). Conservation of T cell receptor conformation in epidermal gammadelta cells with disrupted primary Vgamma gene usage. Science *279*, 1729–1733.

Moran, A.E., Holzapfel, K.L., Xing, Y., Cunningham, N.R., Maltzman, J.S., Punt, J., and Hogquist, K.A. (2011). T cell receptor signal strength in Treg and iNKT cell development demonstrated by a novel fluorescent reporter mouse. J. Exp. Med. *208*, 1279–1289.

Schindelin, J., Arganda-Carreras, I., Frise, E., Kaynig, V., Longair, M., Pietzsch, T., Preibisch, S., Rueden, C., Saalfeld, S., Schmid, B., et al. (2012). Fiji: an open-source platform for biological-image analysis. Nat. Methods 9, 676–682.

Stetson, D.B., Mohrs, M., Reinhardt, R.L., Baron, J.L., Wang, Z.-E., Gapin, L., Kronenberg, M., and Locksley, R.M. (2003). Constitutive cytokine mRNAs mark natural killer (NK) and NK T cells poised for rapid effector function. J. Exp. Med. 198, 1069–1076.

Witherden DA, VerdinoP, Rieder SE, et al. The junctional adhesion moleculeJAML is a costimulatory receptor for. epithelial gammadelta T cell activation. Science. 2010; 329:1205-1210.

Zhang, B., Wu, J., Jiao, Y., Bock, C., Dai, M., Chen, B., Chao, N., Zhang, W., and Zhuang, Y. (2015). Differential requirements of TCR signaling in homeostatic maintenance and function of dendritic epidermal T cells. J. Immunol. *195*, 4282–4291.
